# Broad-range RNA modification analysis of complex biological samples using rapid C18-UPLC-MS

**DOI:** 10.1101/2020.09.03.281246

**Authors:** Pavlina Gregorova, Nina H. Sipari, L. Peter Sarin

## Abstract

Post-transcriptional RNA modifications play an important role in cellular metabolism with homeostatic disturbances manifesting as a wide repertoire of phenotypes, reduced stress tolerance and translational perturbation, developmental defects, and diseases, such as type II diabetes, leukemia and carcinomas. Hence, there has been an intense effort to develop various methods for investigating RNA modifications and their roles in various organisms, including sequencing-based approaches and, more frequently, liquid chromatography mass spectrometry (LC-MS)-based methods. Although LC-MS offers numerous advantages, such as being highly sensitive and quantitative over a broad detection range, some stationary phase chemistries struggle to resolve positional isomers. Furthermore, the demand for detailed analyses of complex biological samples often necessitates long separation times, hampering sample-to-sample turnover and making multisample analyses time consuming. To overcome this limitation, we have developed an ultra-performance LC-MS (UPLC-MS) method that uses an octadecyl carbon chain (C18)-bonded silica matrix for the efficient separation of 50 modified ribonucleosides, including positional isomers, in a single 9 min sample-to-sample run. To validate the performance and versatility of our method, we analyzed tRNA modification patterns of representative microorganisms from each kingdom of life, namely Archaea (*Methanosarcina acetivorans*), Bacteria (*Pseudomonas syringae*) and Eukarya (*Saccharomyces cerevisiae*). Additionally, our method is flexible and readily applicable for detection and relative quantification using stable isotope labelling and targeted approaches like multiple reaction monitoring (MRM). In conclusion, this method represents a fast and robust tool for broad-range exploration and quantification of ribonucleosides, facilitating future homeostasis studies of RNA modification in complex biological samples.

## INTRODUCTION

During the past decade, research into post-transcriptional nucleoside modifications – in particular those of messenger RNA and transfer RNA – has experienced a renaissance as their multifaceted roles, ranging from translational control^1^ via infection^2^ and stress response^3,4^ to development and aging^5,6^, are slowly beginning to unfold. This can be credited to methods and instrument development in two key areas; (I) translation studies using ribosome profiling combined with next-generation sequencing^7,8^, which enables detailed insights into the translational state of cells, and (II) identification and quantitative characterization of modified ribonucleosides by liquid chromatography mass spectrometry (LC-MS). In case of the latter, advances in instrument design, such as improved mass resolution and accuracy, sensitivity, dynamic range, and scan speed, coupled with improved stationary phase chemistries, has expanded the applicability of LC-MS based ribonucleoside modification analysis. However, to achieve a robust and reliable analysis platform with minimal bias, careful consideration must be given to the choice of mobile and stationary phase as well as the solvent delivery system. Currently, reversed-phase high performance liquid chromatography (RP-HPLC) using C18-based materials coupled with electrospray ionization mass spectrometry (ESI-MS) is the prevailing approach used by numerous laboratories^4,9–20^. In addition, bespoke methods have been developed using other stationary phases, such as HILIC^21,22^ and porous graphitic carbon (PGC)^23,24^.

Despite these recent advances, existing HPLC-MS methods are often relatively lengthy as they necessitate separation times ranging from 15 to >70 min, followed by a regeneration step of 5 to 20 min. Consequently, the total sample-to-sample turnover time can be in excess of 90 min, thereby reducing their applicability for analysis of large sample cohorts. To overcome this limitation, several ultra-performance LC-MS (UPLC-MS) methods have been established to accelerate ribonucleoside analyses by reducing overall run times^16,19,22,25^. Nonetheless, these approaches were employed to analyze only a limited number of ribonucleosides. This also restricts their applicability for ribonucleoside analysis as different types of RNA molecules have a significant variability in their modification composition. Therefore, a method suitable for the detection of multiple modifications, especially positional isomers, is desirable. Such an universal method, easily applicable to any organism and type of RNA, would undoubtedly provide a powerful tool for investigation of the crosstalk between the modifications in different RNA molecules^26^, as well as their conservation across the species.

To this end, we initially compared the suitability of three distinct UPLC-compatible stationary phases for broad-range applicability and unbiased separation of modified ribonucleosides. Using a BEH-C18 column, we developed a fast and robust UPLC-MS approach for accurate and quantitative identification of 50 ribonucleosides during a single 9 min analysis run (5 min separation and 4 min column wash and equilibration). Furthermore, we have validated our method by analyzing a complex mix of synthetic standards and biological RNA isolates originating from representative microorganisms of all three kingdoms of life (Archaea, Bacteria and Eukarya). Compared to currently available protocols, the method presented here provides a reduced total runtime and an improved separation of isobaric ribonucleosides, such as positional isomers, which together with unique fingerprint ionization patterns ensures their reliable identification. Taken together, this UPLC-MS method represents a fast, versatile and easily implementable tool for reproducible and reliable ribonucleoside quantification that is readily applicable for large sample cohorts.

## MATERIAL AND METHODS

### Chemicals and ribonucleoside standards

The chemicals used were, unless otherwise stated, purchased from Acros Chemicals and/or Fisher Chemicals in the highest purity level available. The ribonucleoside standards were purchased from Carbosynth (Newbury, UK) and Sigma-Aldrich (canonical ribonucleosides only), including 49 naturally occurring ribonucleosides (full names are in Supplementary Table S1): A, C, G, U, Ψ, A_m_, C_m_, G_m_, U_m_, I, I_m_, m^1^I, m^1^A, m^6^A, m^8^A, m^2,8^A, ms^2^m^6^A, m^6^A_m_, ac^6^A, f^5^C, m^3^C, m^4^C, m^5^C, m^4^C_m_, m^5^C_m_, hm^5^C, m^4,4^C, ac^4^C, ac^4^C_m_, m^1^G, m^2^G, m^7^G, m^2,2^G, m^2,7^G, imG-14, m^3^U, ncm^5^U, m^5^U, m^5^s^2^U, mcm^5^U, mcm^5^s^2^U, cnm^5^U, mo^5^U, cm^5^U, ho^5^U, s^2^C, s^2^U, s^4^U, m^1^Ψ, and 1 non-natural ribonucleoside - m^1,3^Ψ. The purity of each standard is noted in Supplementary Table S1.

### Preparation of standards

Ribonucleoside standards were dissolved in 5 mM ammonium formate (NH_4_HCO_2_), pH 5.3 to concentration of 50 μg/ml. A limited (LM) and complete mix (CM) containing 22 and 50 ribonucleoside standards, respectively, were mixed at concentration of 10 μg/ml. To differentiate the isomeric ribonucleoside modifications, four different mixes (M1-M4) were prepared at final concentration 10 μg/ml. The detailed composition of these mixes is specified in the Supplementary Data (Supplementary Tables S1-S5).

### Biological material, transfer RNA isolation and its preparation for C18-UPLC-MS analysis

*Saccharomyces cerevisiae* BY4741 and *Pseudomonas syringae* pv. *tomato* DC3000 were grown at 28 °C, 180 rpm in 100 ml YPD and King’s B medium, respectively. Cells were harvested by centrifugation at 4 000 *g*, 4 °C, 5 min. *Methanosarcina acetivorans* C2A Δhpt^27^ was cultivated anaerobically at 37 °C, in a single-cell morphology^28^ in 50 ml of high-salt broth medium containing 125 mM methanol (detailed composition in Supplementary Methods). Cultures were harvested anaerobically by centrifugation. The collection time points (t_1_ and t_2_, respectively) were as follows: 1 h and 6 h for *S. cerevisiae*; 1 h and 4 h for *P. syringae*; and 8 h and 27 h for *M. acetivorans*.

The ^15^N metabolically labelled RNA was prepared by growing *P. syringae* in M9 minimal medium supplemented with ^15^NH_4_Cl (Sigma-Aldrich) as the only nitrogen source. Cells were grown O/N and then used for inoculation of new culture. Subsequently, the cells were cultivated until logarithmic growth phase and then were harvested as described above. tRNA was isolated as described below and the labelling efficiency was estimated by MS (data not shown).

Purification of tRNA from *S. cerevisiae* was performed as previously described^18^ with minor changes for *P. syringae* and *M. acetivorans*. Briefly, total RNA was isolated using TRI reagent (Invitrogen) according to manufacturer’s protocol. The RNA containing aqueous phase was transferred to a fresh centrifugation tube and re-isolated by addition of 2 ml acidic phenol and 0.1 vol. of 1-bromo-3-chloropropane (BCP). The suspension was vortexed for 1 min and centrifuged for 15 min at 10 000 *g*. The RNA containing aqueous phase was collected and the volume was adjusted to 10 ml with equilibration buffer EQ (10 mM Tris-HCl, pH 6.3, 15% ethanol, 200 mM KCl), following which it was applied onto a pre-equilibrated (EQ buffer containing 0.15% Triton X-100) Nucleobond AX-100 column (Macherey-Nagel). The column-bound RNA was washed twice with wash buffer WB (10 mM Tris-HCl, pH 6.3, 15% ethanol, 300 mM KCl), and the tRNA containing fraction was eluted with warm (55 °C) elution buffer EB (10 mM Tris-HCl, pH 6.3, 15% ethanol, 750-800 mM KCl) into 2.5 vol. of 99.6% ethanol. The tRNA was precipitated O/N at −20 °C and pelleted by centrifugation at 10 000 *g*, 4 °C for 30 min. Residual salt was washed out with 80% ethanol. The final tRNA pellet was air-dried at room temperature and re-suspended in RNase/DNase-free water. The quality and purity of isolated tRNA was assessed by denaturing electrophoresis on 10% polyacrylamide gel (Supplementary Data). Dephosphorylated mononucleosides for MS analysis were generated as previously described^23^. Prior to MS analysis, the completeness of cleavage was verified by gel electrophoresis. For data normalization, *P. syringae* nucleosides were mixed with ^15^N-labeled ribonucleosides, which served as internal standard (25 ng ^15^N-labeled ribonucleosides per 100 ng of sample).

### UPLC-MS using BEH C18 columns

Unless otherwise stated, all ribonucleoside standards and biological samples were analyzed with Waters Acquity^®^ UPLC system (Waters, Milford MA, USA) attached to Waters Synapt G2 HDMS mass spectrometer (Waters, Milford MA, USA) via an ESI ion source. All samples were analyzed in positive ion mode (sensitivity) with the mass range (m/z) from 100 to 600. The samples were loaded onto the column as follows: single ribonucleoside standards or their mixes, 50 ng; and dephosphorylated mononucleosides originating from biological material, 500 ng. Ribonucleoside separation was performed in Acquity UPLC^®^ Ethylene Bridged Hybrid [BEH] C18 columns (1.7 μm, 2.1 × 50 and 150 mm, Waters, Ireland) at +40 °C. The mobile phase consisted of (A) 0.1% formic acid in MQ H_2_O (Merck Millipore, Bedford, MA, USA) and (B) 0.1% formic acid in acetonitrile (Honeywell, Riedel-de Haën, CHROMASOLV™, LC-MS grade, Steinheim, Germany). Ribonucleoside separation was achieved by a two-step gradient ranging from 1-12% B for 3 min and 12-70% B for another 3 min, followed by a shift to 1% B and subsequent equilibration step for 3 min, with a total run time of 9 min. The flow rate of the mobile phase was 0.45 ml/min, the injection volume was 10 μl and tray temperature was set to 15 °C. Capillary voltage was set to at 3.0 kV, sampling cone 30 and extraction cone 3.0, the source temperature 120 °C and desolvation temperature 400 °C, cone and desolvation gas flow rate was set to 20 l/h and 800 l/h, respectively. Detailed steps of UPLC-MS method development, including ribonucleoside separation on other stationary phases and parameters used for multiple reaction monitoring (MRM) with ABSciex UPLC-MS QTRAP-6500+ instrument are described in Supplementary Methods and Supplementary Table S7.

### Data analysis

Total ion (TIC) and extracted ion (XIC) chromatograms were generated from .RAW format using MZmine2 software (version 2.40.1)^29^ and exported in csv format. Peak identification was performed by custom database identification module of MZmine2 using a lookup list of ribonucleosides consisting of previously reported [M+H]^+^ and product ions masses (data from Modomics^30^); and the retention time information acquired from analyzing available standards. Based on the data obtained with our instrument, this list was further manually curated and additional ions detected in standards and biological samples were added (Figure 2C and Supplementary Figures S3, S6). The final identity assignment was confirmed only when both, [M+H]^+^ and product ions were detected in MS1 spectrum. For data visualization, the relative signal intensities were calculated as a proportion of the most intense signal in the spectrum. To quantify the changes in ribonucleoside modifications, the absolute intensities were first normalized to cytidine and/or to equivalent ^15^N-labeled ribonucleoside (where applicable) to reflect the variances in amount of input tRNA^31^. Subsequently, the relative changes are represented as t_2_/t_1_ ratio of normalized intensities. Error bars in plots represent standard deviations of three independent replicates and p-values were calculated by one-sample Student’s t-test. The data visualization and calculations were done by custom scripts in R (version 3.6.3) using tidyverse package (version 1.3.0)^32^. The calculation of chromatographic parameters for the C18 column is outlined in the Supplementary Methods.

## RESULTS

### C18 UPLC enables short run times and favorable separation and identification of multiple modified ribonucleosides in a single analysis

Ribonucleosides are highly polar compounds and notoriously difficult to analyze on most stationary phases^33^. Hence, we first investigated the elution characteristics of 22 ribonucleosides (LM) on three stationary phases, which are most frequently used for ribonucleoside analysis^4,9,13–16,21–23,34^ – hydrophilic interaction chromatography (HILIC), carbamoyl-coated silica (Amide), and octadecyl carbon chain (C18)-bonded silica – using normal or reversed phase UPLC-compatible setups in 50 or 100 mm column format (Supplementary Figure S1). The initial experiments using our UPLC-MS conditions showed that separation of LM standards on BEH-HILIC and BEH-AMIDE columns (both 100 mm long) was insufficient, especially for the positional isomers. Additionally, we also observed substantial peak tailing and unsatisfactory signal-to-noise ratios (Supplementary Figure S1A-D). On the other hand, the 50 mm BEH-C18 column was able to achieve the fastest separation time (<3 min), while providing more favorable signal-to-noise ratios and an exceptional separation for all tested ribonucleosides, including positional isomers (Supplementary Figure S1E). Thus, we continued the method development using only this stationary phase.

To obtain a method by which multiple modified ribonucleosides can be identified in a single analysis, we increased the number of ribonucleoside standards to 50 (see Experimental section) and assessed their individual elution and ionization characteristics (Supplementary Figure S2 and S3), as well as their chromatographic parameters (Supplementary Table S6). Next, we decided to increase the challenge by analyzing these 50 standards in a complex mix (CM) using our UPLC-MS coupled to the BEH-C18 column. However, the 50 mm BEH-C18 column proved inadequate for reliable identification of all the CM standards (data not shown). Therefore, we examined the CM separation on a longer 150 mm column with the same BEH-C18 stationary phase (Figure 1A). To facilitate the identification of all available isomers, we prepared four mixes of standards (M1-4), where each mix contained only one of the available isobaric isomers (Figure 1B, Supplementary Figure S4). The examination of M1-4 XICs overlays showed that most isobaric isomers achieve baseline separation (Figure 1C, Supplementary Figure S5A). Nevertheless, some positional isomers including methylated cytidines, methylated inosines, and methylated guanosines, eluted too closely to each other and prevented their differentiation solely by retention time (Figure 1C-D). Therefore, we closely re-examined the individual spectra of these isomers (Supplementary Figure S3) and compared their fragmentation patterns. In the case of methylated inosines (Figure 1D, right panel), the spectral comparison provided a confident measure for isomer identification. Similarly, this approach also proved sufficient for distinction of G_m_ and m^2^G/m^1^G, which elute closely to each other (Supplementary Figure S5B). However, this approach was not applicable for distinguishing base methylated cytidines.

**Figure 1.**
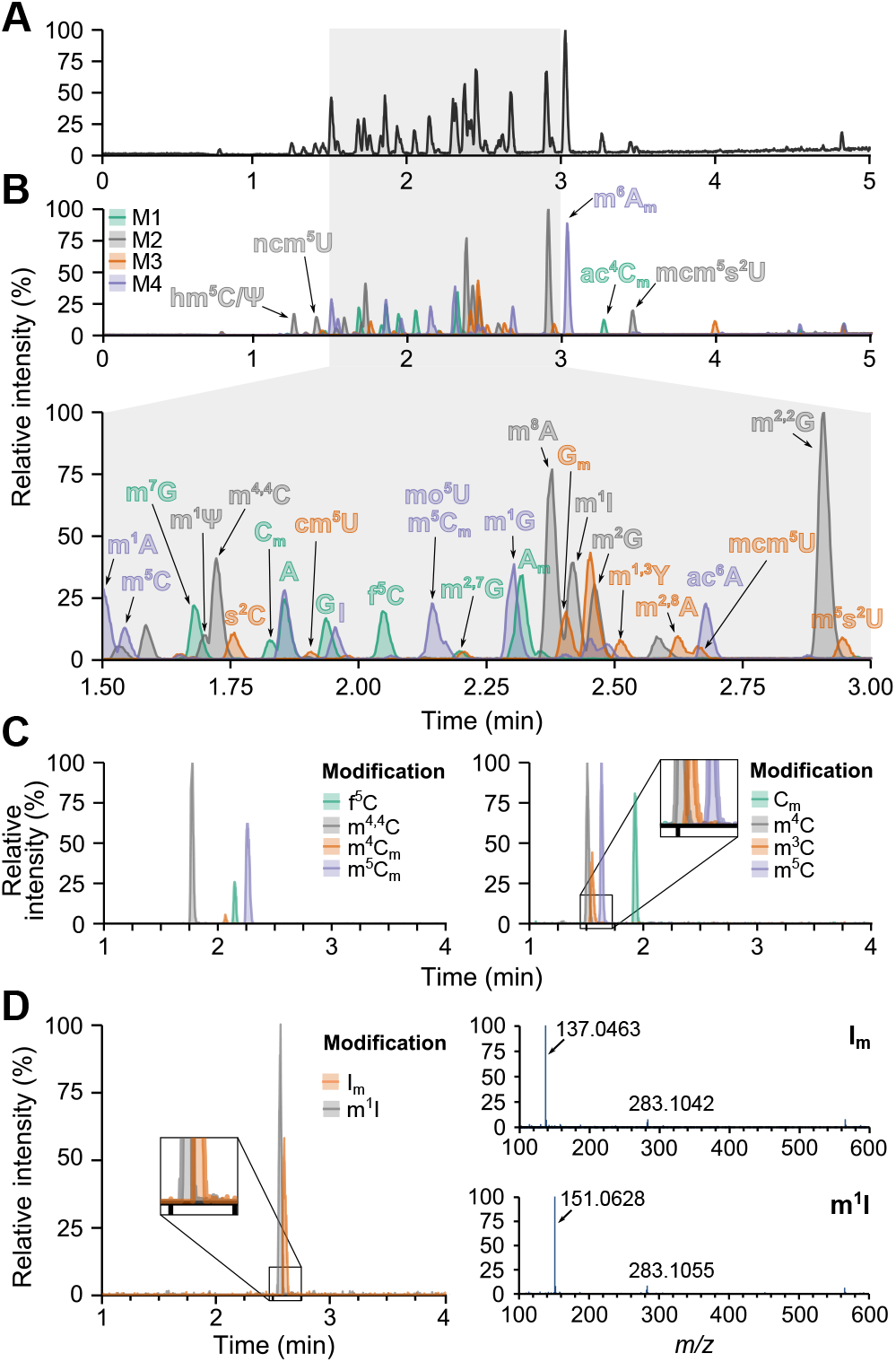
C18 UPLC-MS provides baseline separation of 50 ribonucleosides and enables their reliable identification. **(A)** Total ion chromatogram (TIC) of CM of standards analyzed on 150 mm C18 column and **(B)** overlaid TICs of M1-M4. **(C)** The separation of selected isobaric and positional isomers on 150 mm C18 column represented as extracted ion chromatograms (XICs extracted from C18 UPLC-MS analysis of M1-M4. **(D)** XICs of selected positional isomers (left panel) and their respective MS spectra (right panel) showing the distinct ionization profile.

The comparison of our recorded spectra to previously published data^30^ revealed an additional variation in the fragmentation patterns (Supplementary Figure S3, in red), namely the addition of dimer ions ([2M+H]+). Dimerization occurs in the ESI chamber and is influenced by the LC-conditions (pH, solvents) and ES ionization parameters (voltage, temperature etc.), as well as the chemical properties and concentration of the analytes. Since the analysis conditions constitute the best possible compromise for all 50 analytes, dimer formation (Supplementary Figure S3) cannot be entirely avoided without experiencing a significant reduction in chromatographic resolution or ionization efficiency. In the case of specific analytes, we observed dimers in the ribonucleoside standards throughout a wide concentration range as well as in the biological samples. This dimerization pattern proved beneficial in assigning the identity of co-eluting positional isomers. For example, the spectra of methylated cytidines m^4^C, m^5^C and C_m_ contains an ion at *m/z* 515, whereas this ion is absent in m^3^C (Supplementary Figure S5C). Similar fingerprint ionization patterns for distinction of isomers has been characterized before using higher-energy collisional dissociation MS^35^.

### C18 UPLC-MS provides robust identification of ribonucleosides in biological samples from various organisms

While CM standards resemble the complexity of biological samples, the RNA isolates often contain impurities that may impair MS analysis. Transfer RNA (tRNA) contains on average 13 ribonucleoside modifications per molecule, although the exact number and type of modifications vary between different organisms and tissue types or due to growth conditions and/or presence of stress^3,4,8,30,36^. To determine if our method can robustly identify and quantitatively characterize relative modification changes, we decided to analyze ribonucleoside modifications in total tRNA of three representative microorganisms – *M. acetivorans* (Archaea), *P. syringae* (Bacteria) and *S. cerevisiae* (Eukarya) (Figure 2A, Supplementary Figure S6). In agreement with previously published data^30,37^, we detected a total of 32 tRNA modifications, including anticodon modifications, such as mcm^5^U_34_, mcm^5^s^2^U_34_ and m^1^I_37_ in Eukarya^38–40^ and s^2^C_32_ in Bacteria^41^ (Figure 2B). Next, we compared the detected modifications between the organisms included in the study, highlighting the presence of both universal and specific modifications, which are present only in a particular kingdom of life. For example, s^2^C and m^3^U can be found only in Bacteria, whereas imG-14 is an archaeal modification; mcm^5^U, mcm^5^s^2^U, and m^1^I are eukaryotic modifications, and m^1^G, m^2^G, m^6^A, Ψ, C_m_ are universally conserved throughout all organisms (Figure 2B). On the other hand, rRNA-specific modifications^30,37^, such as m^8^A, m^2,8^A, and m^4^C, were not observed, indicating a sufficient purity of our tRNA isolates (Supplementary Figure S6A). Upon a detailed analysis of the data, we observed additional peaks that were unassigned, as they did not match any of the available standards. To identify these peaks, we created a custom lookup list, which allowed identification by matching their spectra with published data^30^. The confidence of identification was further increased by considering only those modifications for which all previously published ions were detected. Using this approach and the known presence of particular modifications in tRNAs across kingdoms of life, we were able to identify the following modifications: archaeosine (G+), agmatidine (C+) in Archaea; epoxyqueuosine (oQ), 2-methyladenosine (m^2^A) and *N6*-(cis-hydroxyisopentenyl) adenosine (io^6^A) in Bacteria; *N6*-isopentenyladenosine (i^6^A) in Bacteria and Eukarya, and *N6*-threonyl-carbamoyladenosine (t^6^A) in all organisms^30,42,43^ (Figure 2B-C, Supplementary Figure S6B and S6C). As for the unassigned fifth methylated adenosine peak, we utilized the retention times of the known positional isomers (A_m_, m^8^A, m^1^A and m^6^A), spectral matching and its occurrence, which is limited to bacterial samples, to conclude that its identity is most likely that of m^2^A (Figure 2C).

**Figure 2.**
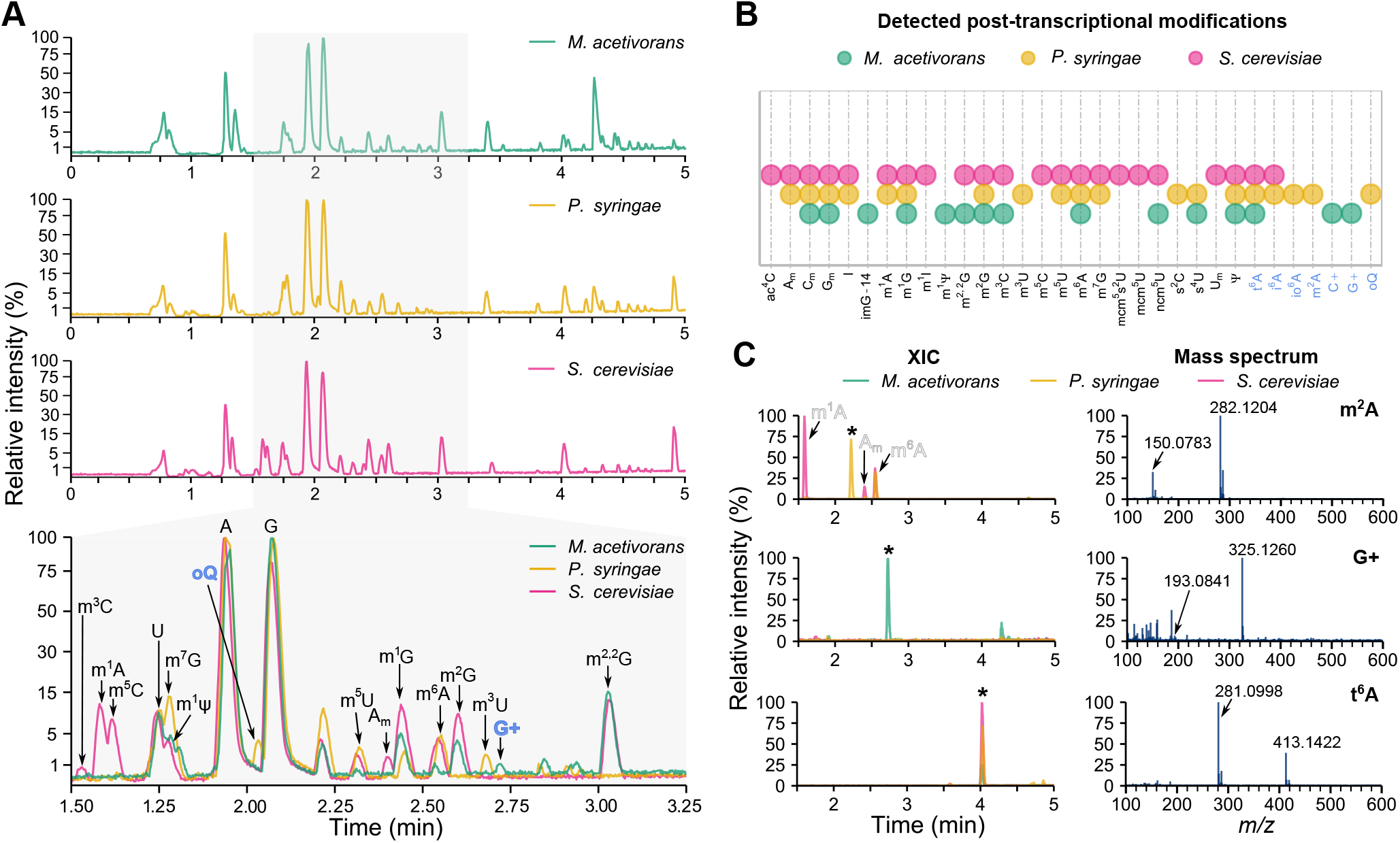
Analysis of tRNA modifications in representative microorganisms of each kingdom of life. **(A)** Individual TICs of archaeal (*M. acetivorans*), bacterial (*P. syringae*) and eukaryotic (*S. cerevisiae*) modifications and their overlay showing the variability of tRNA modifications across kingdoms of life. Zoomed section displays the differences in modifications present in each organism and modifications identified without standards are in blue. **(B)** Summary of detected modifications in each microorganism. Modifications identified without standards are in blue. **(C)** Identification of unknown peaks present in biological samples. Overlay of XICs across all three organisms (left panel) and their respective mass spectra (right panel) highlighting the unique ionization profiles. Arrows in spectra point to neutral loss ions. In XICs, the peak of interest is marked by *.

High-resolution baseline separation of individual analytes requires high stability of the stationary phase over multiple analyses. To determine the suitability of the BEH-C18 column for large-scale studies, we analyzed the retention time variance for the canonical nucleosides throughout 80 consecutive runs (Figure 3A-B). The observed shifts in retention times were negligible, achieving a mean variance of 0.05-0.06 min (Figure 3B), thus confirming the high reproducibility of the method and its suitability for studying large sample cohorts.

**Figure 3.**
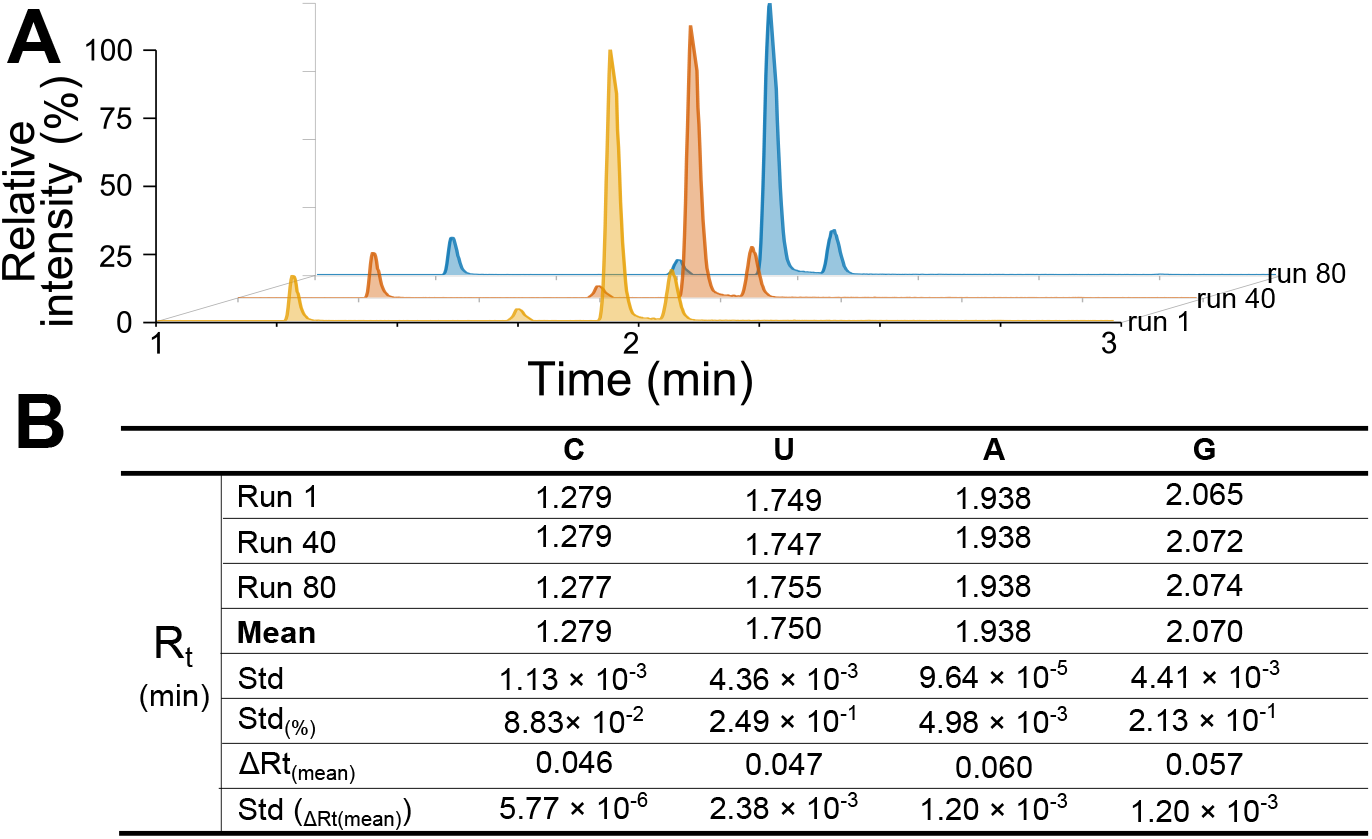
C18 UPLC-MS provides robust and reproducible quantification of modified ribonucleosides. **(A)** Three representative samples selected from consecutive runs showing the XICs of the canonical ribonucleosides. **(B)** Retention time variability over multiple runs. Rt represents retention time at apex of each peak, std equals standard deviation and ΔRt(mean) is the mean difference between the start and the end of peak.

### Relative quantification of ribonucleosides by C18 UPLC-MS

As cells grow, nutrients become increasingly scarce and potentially toxic metabolic byproducts become more abundant, creating an imbalance that causes nutritional stress. Biotic stress is reflected in tRNA modification levels as protein synthesis must adapt to maintain cellular homeostasis^44–48^. Therefore, quantifying tRNA modification changes provides insights that allow us to understand how protein synthesis is modulated by post-transcriptional modifications. To investigate the suitability of our method for such studies, we analyzed the changes in tRNA modification levels of cells harvested at the early exponential growth (t_1_) and late exponential growth phases (t_2_) in set of three independent replicates (n=3) (Figure 4).

**Figure 4.**
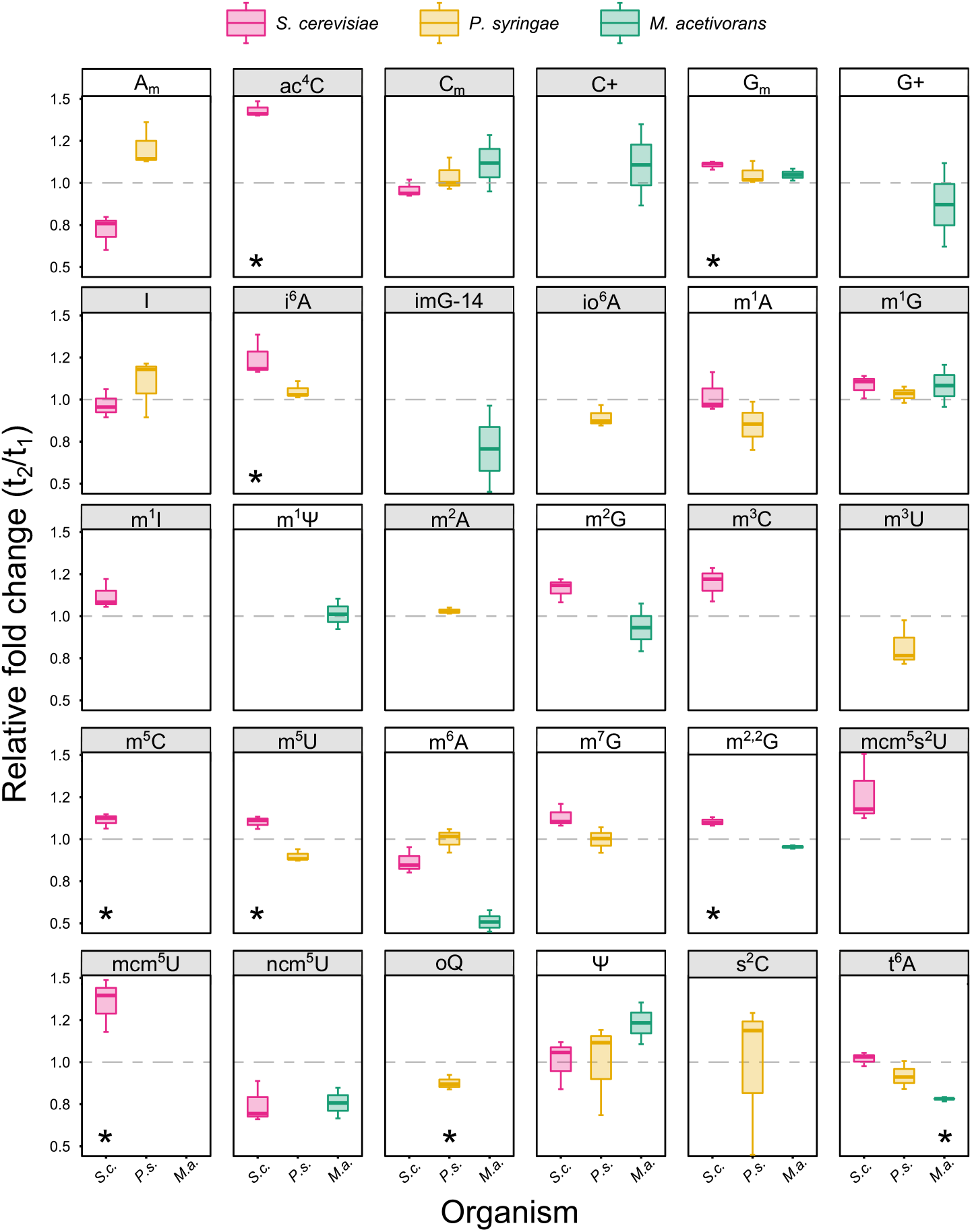
Relative quantification of tRNA modification changes between early exponential growth (t_1_) and late exponential growth phases (t_2_). S.c. - *S. cerevisiae*, P.s. - P. *syringae*, M.a. - *M. acetivorans*. Panels in grey show anticodon stem loop modifications. Grey line highlights value 1 (no change). Statistical significance was calculated using one-sample Student’s t-test and * shows values with p < 0.05 (n = 3).

First, we utilized internal normalization with metabolically labeled ribonucleosides^17,23^ by preparing a batch of [^15^N] stable isotope labeled tRNA of *P. syringae* and analyzing it as a spike-in mixture with non-labeled ribonucleosides (see Material and Methods). Our analysis showed that [^15^N]-labeled ribonucleosides can be easily detected for a majority of all ribonucleosides present in *P. syringae* (Supplementary Figure S7). This approach significantly improves the accuracy of relative quantification by eliminating the variance that stems from the differences in ribonucleoside ionization efficiency^23^. However, metabolic labeling is not without its limitations. To cover all ribonucleosides, spike-in batches must originate from the same organism as the non-labeled sample and, while this may be relatively straightforward and cost-efficient for microbes, it becomes a significant limiting factor for many other model organisms, not to mention tissues and clinical samples.

Consequently, we decided to normalize the peak intensities to the cytidine signal to mitigate intra-sample loading variances^31^. This approach also allowed a comparison of the relative modification changes between all tested organisms (Figure 4). Unsurprisingly, we observed significant changes in abundance for several anticodon stem loop modifications, such as i^6^A, t^6^A, mcm^5^U, mcm^5^s^2^U, and ncm^5^U (Figure 4, grey panels). These modifications alter the processivity and fidelity of translation and for example, mcm^5^s^2^U_34_ has been shown to be critical for the efficient translation of certain stress-induced gene transcripts^20,49,50^. By optimizing conditions for multiple reaction monitoring mode (MRM) with QQQ detector (parameters in Supplementary Table S7), we further increased the detection sensitivity, which revealed modifications undetectable by QTOF. Consequently, our C18 UPLC-MS approach offers a flexible method suitable for both broad-range and targeted characterization as well as relative quantification of tRNA modifications and their changes in response to environmental conditions.

## DISCUSSION

Here, we present an improved and fast UPLC-MS method for the simultaneous identification of 50 ribonucleosides. Thanks to the utilization of UPLC separation on BEH-C18 column, we achieved short ribonucleoside separation (<5 min) and column regeneration times, resulting in a total analysis time of only 9 minutes. This approach provides reproducible and reliable relative quantification of global nucleoside modification changes in various organisms. Furthermore, we also show that our method is directly compatible with previously published spike-in [^15^N] stable isotope labeling techniques for signal normalization^17,23^, which further increase the quantification accuracy. This UPLC-MS method can also be used for dynamic monitoring of tRNA modification changes via pulse-chase labelling approaches, such as NAIL-MS^16^.

In this study, we have validated our method using nucleoside standards (Figure 1) and tRNA (Figure 2), which features the highest diversity of post-transcriptional nucleoside modification of any RNA species. Importantly, our method is highly flexible and due to the number of identified modifications, it can be easily implemented for the complete modification, i.e. ‘modificome’, analysis of other RNA species, such as ribosomal RNA and messenger RNA. However, due to a significantly lower proportion of modified ribonucleosides in these RNA species, the use of more sensitive MS detectors, such as QQQ, or enrichment of target molecules prior analysis might be necessary. In addition, we have shown that in some cases nucleoside modifications can be identified and quantified even without standards by matching the ionization profiles to previously published data. Nevertheless, the lack of standards does limit the applicability of the method for absolute quantification, as this necessitates the use of ribonucleoside-specific calibration curves for the instrument. Future developments will focus on analyzing additional standards and utilizing tandem mass spectrometry to confirm the identity of those modifications, for which standards are currently unavailable. In conclusion, our UPLC-MS method constitutes the first robust and universally applicable tool for fast detection and quantification of RNA modifications in complex biological samples.

## ACKNOWLEDGEMENT

Dr Elina Roine and Prof. Sebastian Leidel are thanked for providing the *P. syringae* and *S. cerevisiae* strain, respectively, and Assistant Prof. Silvan Scheller’s lab is thanked for providing *M. acetivorans* cultures. The authors also wish to thank Dr Ulrike Abendroth for help with RNA isolation and all RNAcious lab members for their comments and discussions.

## REFERENCES

1. Roundtree IA, Evans ME, Pan T, He C. Dynamic RNA Modifications in Gene Expression Regulation. Cell 2017; 169:1187–200. PMID:28622506; https://doi.org/10.1016/j.cell.2017.05.045

2. Koh CS, Sarin LP. Transfer RNA modification and infection –Implications for pathogenicity and host responses. Biochim Biophys Acta - Gene Regul Mech 2018; 1861:419–32. PMID:29378328; https://doi.org/10.1016/j.bbagrm.2018.01.015

3. Huber SM, Leonardi A, Dedon PC, Begley TJ. The Versatile Roles of the tRNA Epitranscriptome during Cellular Responses to Toxic Exposures and Environmental Stress. Toxics 2019; 7:17. PMID:30934574; https://doi.org/10.3390/toxics7010017

4. Chan CTY, Dyavaiah M, DeMott MS, Taghizadeh K, Dedon PC, Begley TJ. A Quantitative Systems Approach Reveals Dynamic Control of tRNA Modifications during Cellular Stress. PLoS Genet 2010; 6:e1001247. PMID:21187895; https://doi.org/10.1371/journal.pgen.1001247

5. Blanco S, Frye M. Role of RNA methyltransferases in tissue renewal and pathology. Curr Opin Cell Biol 2014; 31:1–7. PMID:25014650; https://doi.org/10.1016/j.ceb.2014.06.006

6. Min, K-W, Zealy RW, Davila S, Fomin M, Cummings JC, Makowsky D, Mcdowell CH, Thigpen H, Hafner M, Kwon, S-H, et al. Profiling of m^6^A RNA modifications identified an age-associated regulation of AGO2 mRNA stability. Aging Cell 2018; 17:e12753–e12753. PMID:29573145; https://doi.org/10.1111/acel.12753

7. Ingolia NT, Brar GA, Rouskin S, McGeachy AM, Weissman JS. The ribosome profiling strategy for monitoring translation in vivo by deep sequencing of ribosome-protected mRNA fragments. Nat Protoc 2012; 7:1534–50. PMID:22836135; https://doi.org/10.1038/nprot.2012.086

8. Chou, H-JJ, Donnard E, Gustafsson HT, Garber M, Rando OJ. Transcriptome-wide Analysis of Roles for tRNA Modifications in Translational Regulation. Mol Cell 2017; 68:978–992.e4. PMID:29198561; https://doi.org/10.1016/j.molcel.2017.11.002

9. Weimann A, Belling D, Poulsen HE. Quantification of 8-oxo-guanine and guanine as the nucleobase, nucleoside and deoxynucleoside forms in human urine by high-performance liquid chromatography–electrospray tandem mass spectrometry. Nucleic Acids Res 2002; 30:e7–e7. PMID:11788733; https://doi.org/10.1093/nar/30.2.e7

10. Thüring K, Schmid K, Keller P, Helm M. Analysis of RNA modifications by liquid chromatography–tandem mass spectrometry. Methods 2016; 107:48–56. https://doi.org/10.1016/J.YMETH.2016.03.019

11. Yan M, Wang Y, Hu Y, Feng Y, Dai C, Wu J, Wu D, Zhang F, Zhai Q. A high-throughput quantitative approach reveals more small RNA modifications in mouse liver and their correlation with diabetes. Anal Chem 2013; 85:12173–81. PMID:24261999; https://doi.org/10.1021/ac4036026

12. Zhu Y, Zhou G, Yu X, Xu Q, Wang K, Xie D, Yang Q, Wang L. LC-MS-MS quantitative analysis reveals the association between FTO and DNA methylation. PLoS One 2017; 12:e0175849.

13. Su D, Chan CTY, Gu C, Lim KS, Chionh YH, McBee ME, Russell BS, Babu IR, Begley TJ, Dedon PC. Quantitative analysis of ribonucleoside modifications in tRNA by HPLC-coupled mass spectrometry. Nat Protoc 2014; 9:828–41. PMID:24625781; https://doi.org/10.1038/nprot.2014.047

14. Russell SP, Limbach PA. Evaluating the reproducibility of quantifying modified nucleosides from ribonucleic acids by LC-UV-MS. J Chromatogr B Anal Technol Biomed Life Sci 2013; 923–924:74–82. https://doi.org/10.1016/j.jchromb.2013.02.010

15. Kang B Il, Miyauchi K, Matuszewski M, D’Almeida GS, Rubio MAT, Alfonzo JD, Inoue K, Sakaguchi Y, Suzuki TT, Sochacka E, et al. Identification of 2-methylthio cyclic N6-threonylcarbamoyladenosine (ms^2^ct^6^A) as a novel RNA modification at position 37 of tRNAs. Nucleic Acids Res 2016; 45:2124–36. PMID:27913733; https://doi.org/10.1093/nar/gkw1120

16. Reichle VF, Kaiser S, Heiss M, Hagelskamp F, Borland K, Kellner S. Surpassing limits of static RNA modification analysis with dynamic NAIL-MS. Methods 2019; 156:91–101. PMID:30395967; https://doi.org/https://doi.org/10.1016/j.ymeth.2018.10.025

17. Kellner S, Neumann J, Rosenkranz D, Lebedeva S, Ketting RF, Zischler H, Schneider D, Helm M. Profiling of RNA modifications by multiplexed stable isotope labelling. Chem Commun 2014; 50:3516–8. PMID:24567952; https://doi.org/10.1039/C3CC49114E

18. Alings F, Sarin LP, Fufezan C, Drexler HCA, Leidel SA. An evolutionary approach uncovers a diverse response of tRNA 2-thiolation to elevated temperatures in yeast. RNA 2015; 21:202–12. PMID:25505025; https://doi.org/10.1261/rna.048199.114

19. Basanta-Sanchez M, Temple S, Ansari SA, D’Amico A, Agris PF. Attomole quantification and global profile of RNA modifications: Epitranscriptome of human neural stem cells. Nucleic Acids Res 2015; 44:e26–e26. https://doi.org/10.1093/nar/gkv971

20. Deng W, Babu IR, Su D, Yin S, Begley TJ, Dedon PC. Trm9-Catalyzed tRNA Modifications Regulate Global Protein Expression by Codon-Biased Translation. PLOS Genet 2015; 11:e1005706.

21. Sakaguchi Y, Miyauchi K, Kang B Il, Suzuki T. Nucleoside Analysis by Hydrophilic Interaction Liquid Chromatography Coupled with Mass Spectrometry. In: Methods in Enzymology. Academic Press; page 19–28. https://doi.org/10.1016/bs.mie.2015.03.015

22. Zhou G, Pang H, Tang Y, Yao X, Ding Y, Zhu S, Guo S, Qian D, Shen J, Qian Y, et al. Hydrophilic interaction ultra-performance liquid chromatography coupled with triple-quadrupole tandem mass spectrometry (HILIC-UPLC-TQ-MS/MS) in multiple-reaction monitoring (MRM) for the determination of nucleobases and nucleosides in ginkgo seeds. Food Chem 2014; 150:260–6. PMID:24360448; https://doi.org/10.1016/j.foodchem.2013.10.143

23. Sarin LP, Kienast SD, Leufken J, Ross RL, Dziergowska A, Debiec K, Sochacka E, Limbach PA, Fufezan C, Drexler HCA, et al. Nano LC-MS using capillary columns enables accurate quantification of modified ribonucleosides at low femtomol levels. RNA 2018; 24:1403–17. PMID:30012570; https://doi.org/10.1261/rna.065482.117

24. Giessing AMB, Scott LG, Kirpekar F. A Nano-Chip-LC/MSn Based Strategy for Characterization of Modified Nucleosides Using Reduced Porous Graphitic Carbon as a Stationary Phase. J Am Soc Mass Spectrom 2011; 22. https://doi.org/10.1021/jasms.8b04060

25. Šimonová A, Svojanovská B, Trylčová J, Hubálek M, Moravčík O, Zavřel M, Pávová M, Hodek J, Weber J, Cvačka J, et al. LC/MS analysis and deep sequencing reveal the accurate RNA composition in the HIV-1 virion. Sci Rep 2019; 9:8697. PMID:31213632; https://doi.org/10.1038/s41598-019-45079-1

26. Ontiveros RJ, Shen H, Stoute J, Yanas A, Cui Y, Zhang Y, Liu KF. Coordination of mRNA and tRNA methylations by TRMT10A. Proc Natl Acad Sci 2020; 117:7782–91. PMID:32213595; https://doi.org/10.1073/PNAS.1913448117

27. Pritchett MA, Zhang JK, Metcalf WW. Development of a markerless genetic exchange method for Methanosarcina acetivorans C2A and its use in construction of new genetic tools for methanogenic archaea. Appl Environ Microbiol 2004; 70:1425–33. PMID:15006762; https://doi.org/10.1128/aem.70.3.1425-1433.2004

28. Sowers KR, Boone JE, Gunsalus RP. Disaggregation of Methanosarcina spp. and Growth as Single Cells at Elevated Osmolarity. Appl Environ Microbiol 1993; 59:3832–9. PMID:16349092;

29. Pluskal T, Castillo S, Villar-Briones A, Orešič M. MZmine 2: Modular framework for processing, visualizing, and analyzing mass spectrometry-based molecular profile data. BMC Bioinformatics 2010; 11:395. PMID:20650010; https://doi.org/10.1186/1471-2105-11-395

30. Boccaletto P, Machnicka MA, Purta E, Piatkowski P, Baginski B, Wirecki TK, de Crécy-Lagard V, Ross R, Limbach PA, Kotter A, et al. MODOMICS: a database of RNA modification pathways. 2017 update. Nucleic Acids Res 2018; 46:D303–7. PMID:29106616; https://doi.org/10.1093/nar/gkx1030

31. Cai WM, Chionh YH, Hia F, Gu C, Kellner S, McBee ME, Ng CS, Pang YLJ, Prestwich EG, Lim KS, et al. A Platform for Discovery and Quantification of Modified Ribonucleosides in RNA: Application to Stress-Induced Reprogramming of tRNA Modifications. Methods Enzymol 2015; 560:29–71. https://doi.org/10.1016/bs.mie.2015.03.004

32. Wickham H, Averick M, Bryan J, Chang W, McGowan L, François R, Grolemund G, Hayes A, Henry L, Hester J, et al. Welcome to the Tidyverse. J Open Source Softw 2019; 4:1686. https://doi.org/10.21105/joss.01686

33. Gehrke CW, Kuo KC. Ribonucleoside analysis by reversed-phase high-performance liquid chromatography. J Chromatogr 1989; 471:3–36. PMID:2670985; https://doi.org/10.1016/s0021-9673(00)94152-9

34. Kellner S, Ochel A, Thüring K, Spenkuch F, Neumann J, Sharma S, Entian K-D, Schneider D, Helm M. Absolute and relative quantification of RNA modifications via biosynthetic isotopomers. Nucleic Acids Res 2014; 42:e142–e142. PMID:25129236; https://doi.org/10.1093/nar/gku733

35. Jora M, Burns AP, Ross RL, Lobue PA, Zhao R, Palumbo CM, Beal PA, Addepalli B, Limbach PA. Differentiating Positional Isomers of Nucleoside Modifications by Higher-Energy Collisional Dissociation Mass Spectrometry (HCD MS). J Am Soc Mass Spectrom 2018; 29:1745–56. PMID:29949056; https://doi.org/10.1007/s13361-018-1999-6

36. Pan T. Modifications and functional genomics of human transfer RNA. Cell Res 2018; 28:395–404. PMID:29463900; https://doi.org/10.1038/s41422-018-0013-y

37. Cantara WA, Crain PF, Rozenski J, McCloskey JA, Harris KA, Zhang X, Vendeix FAP, Fabris D, Agris PF. The RNA modification database, RNAMDB: 2011 update. Nucleic Acids Res 2010; 39:D195–201. PMID:21071406; https://doi.org/10.1093/nar/gkq1028

38. Nedialkova DD, Leidel SA. Optimization of Codon Translation Rates via tRNA Modifications Maintains Proteome Integrity. Cell 2015; 161:1606–18. PMID:26052047; https://doi.org/10.1016/j.cell.2015.05.022

39. Hou Y-M, Perona JJ. Stereochemical mechanisms of tRNA methyltransferases. FEBS Lett 2010; 584:278–86. PMID:19944101; https://doi.org/10.1016/j.febslet.2009.11.075

40. Karlsborn T, Tükenmez H, Mahmud AKMF, Xu F, Xu H, Byström AS. Elongator, a conserved complex required for wobble uridine modifications in eukaryotes. RNA Biol 2014; 11:1519–28. PMID:25607684; https://doi.org/10.4161/15476286.2014.992276

41. Jäger G, Leipuviene R, Pollard MG, Qian Q, Björk GR. The conserved Cys-X1-X2-Cys motif present in the TtcA protein is required for the thiolation of cytidine in position 32 of tRNA from Salmonella enterica serovar Typhimurium. J Bacteriol 2004; 186:750–7. PMID:14729701; https://doi.org/10.1128/jb.186.3.750-757.2004

42. Hori H. Methylated nucleosides in tRNA and tRNA methyltransferases. Front Genet 2014; 5:144. PMID:24904644; https://doi.org/10.3389/fgene.2014.00144

43. Schweizer U, Bohleber S, Fradejas-Villar N. The modified base isopentenyladenosine and its derivatives in tRNA. RNA Biol 2017; 14:1197–208. PMID:28277934; https://doi.org/10.1080/15476286.2017.1294309

44. Barraud P, Tisné C. To be or not to be modified: Miscellaneous aspects influencing nucleotide modifications in tRNAs. IUBMB Life 2019; 71:1126–40. PMID:30932315; https://doi.org/10.1002/iub.2041

45. Chan CTY, Pang YLJ, Deng W, Babu IR, Dyavaiah M, Begley TJ, Dedon PC. Reprogramming of tRNA modifications controls the oxidative stress response by codon-biased translation of proteins. Nat Commun 2012; 3:937. https://doi.org/10.1038/ncomms1938

46. Tuorto F, Legrand C, Cirzi C, Federico G, Liebers R, Müller M, Ehrenhofer-Murray AE, Dittmar G, Gröne H, Lyko F. Queuosine-modified tRNAs confer nutritional control of protein translation. EMBO J 2018; 37. PMID:30093495; https://doi.org/10.15252/embj.201899777

47. Zaborske JM, DuMont VL, Wallace EW, Pan T, Aquadro CF, Drummond DA. A nutrient-driven tRNA modification alters translational fidelity and genome-wide protein coding across an animal genus. PLoS Biol 2014; 12:e1002015. PMID:25489848; https://doi.org/10.1371/journal.pbio.1002015

48. Zaborske JM, Bauer DuMont VL, Wallace EWJ, Pan T, Aquadro CF, Drummond DA. Correction: A nutrient-driven tRNA modification alters translational fidelity and genome-wide protein coding across an animal genus. PLoS Biol 2015; 13:e1002150–e1002150. https://doi.org/10.1371/journal.pbio.1002150

49. Rezgui VA, Tyagi K, Ranjan N, Konevega AL, Mittelstaet J, Rodnina M V, Peter M, Pedrioli PG. tRNA tKUUU, tQUUG, and tEUUC wobble position modifications fine-tune protein translation by promoting ribosome A-site binding. Proc Natl Acad Sci U S A 2013; 110:12289–94. PMID:23836657; https://doi.org/10.1073/pnas.1300781110

50. Bauer F, Matsuyama A, Candiracci J, Dieu M, Scheliga J, Wolf DA, Yoshida M, Hermand D. Translational control of cell division by Elongator. Cell Rep 2012; 1:424–33. https://doi.org/10.1016/j.celrep.2012.04.001

