## Supplementary Data for "Broad-range RNA modification analysis of complex biological samples using rapid C18-UPLC-MS"

**SUPPLEMENTARY FIGURE S1**

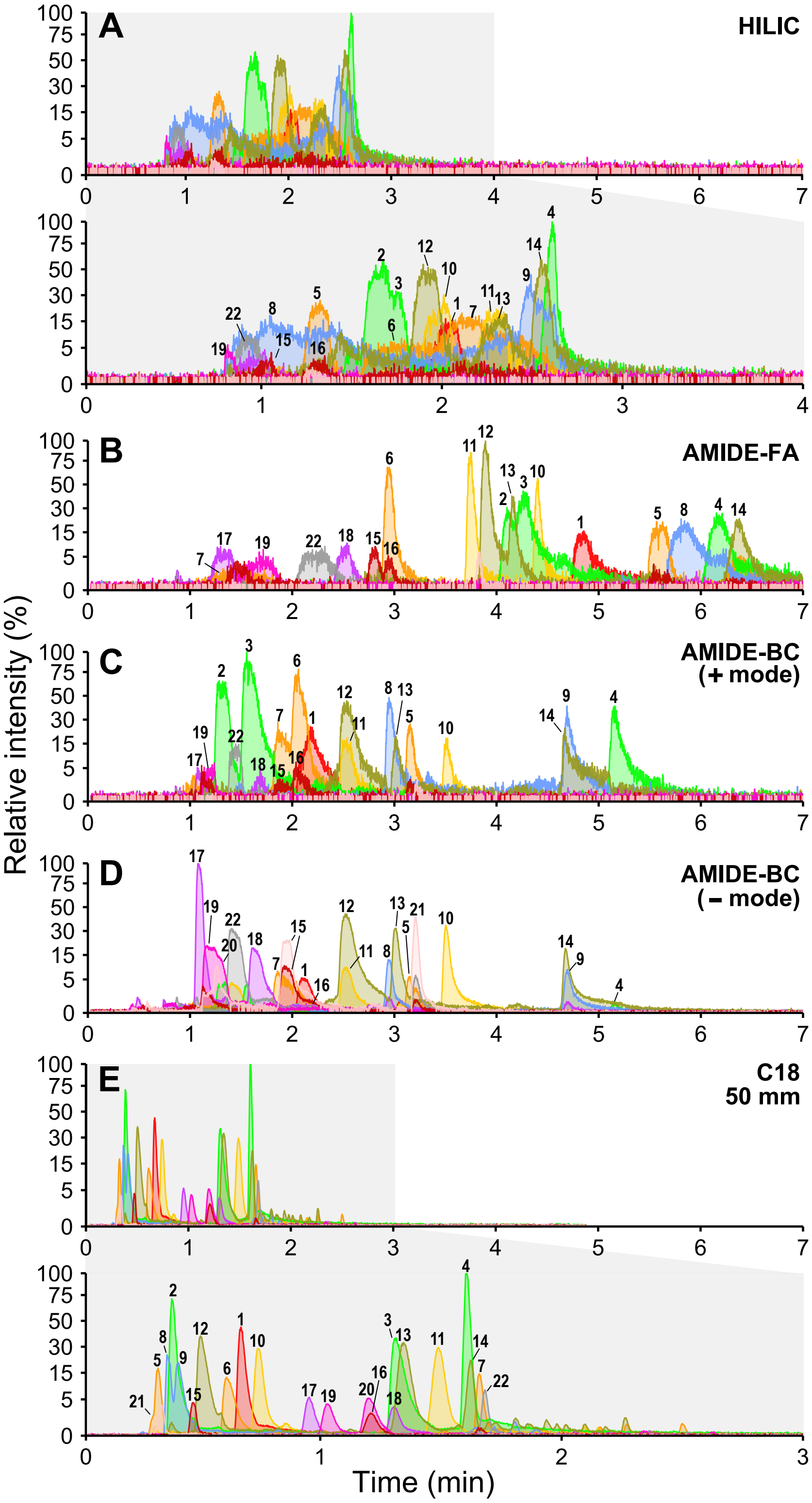


**Supplementary Figure S1.** Extracted chromatograms (XICs) of limited mix (LM) of nucleoside standards showing the LC separation profiles of different stationary phases. **(A)** HILIC with 0.1% formic acid (FA) as solvent A, **(B-D)** AMIDE with two different solvents A - 0.1% FA and 10 mM ammonium bicarbonate, pH 9 (BC) in both positive (+) and negative (-) mode. **(E)** C18 column (50 mm) with 0.1% formic acid as solvent A. Acetonitrile (ACN) was used as solvent B in all runs. The peak legend is as follows: 1 – A, 2 – m$^1$A, 3 – A$_m$, 4 – m$^6$A, 5 – C, 6 – C$_m$, 7 – ac$^4$C, 8 – m$^3$C, 9 – m$^5$C, 10 – G, 11 - G$_m$, 12 – m$^7$G, 13 – m$^2$G, 14 – m$^1$G, 15 - U, 16 - U$_m$, 17 – m$^5$U, 18 – m$^3$U, 19 – s$^2$U, 20 – s$^4$U, 21 – Ψ, 22 – mcm$^5$U. Isomeric modified ribonucleosides are depicted in the same color.

#### SUPPLEMENTARY FIGURE S2

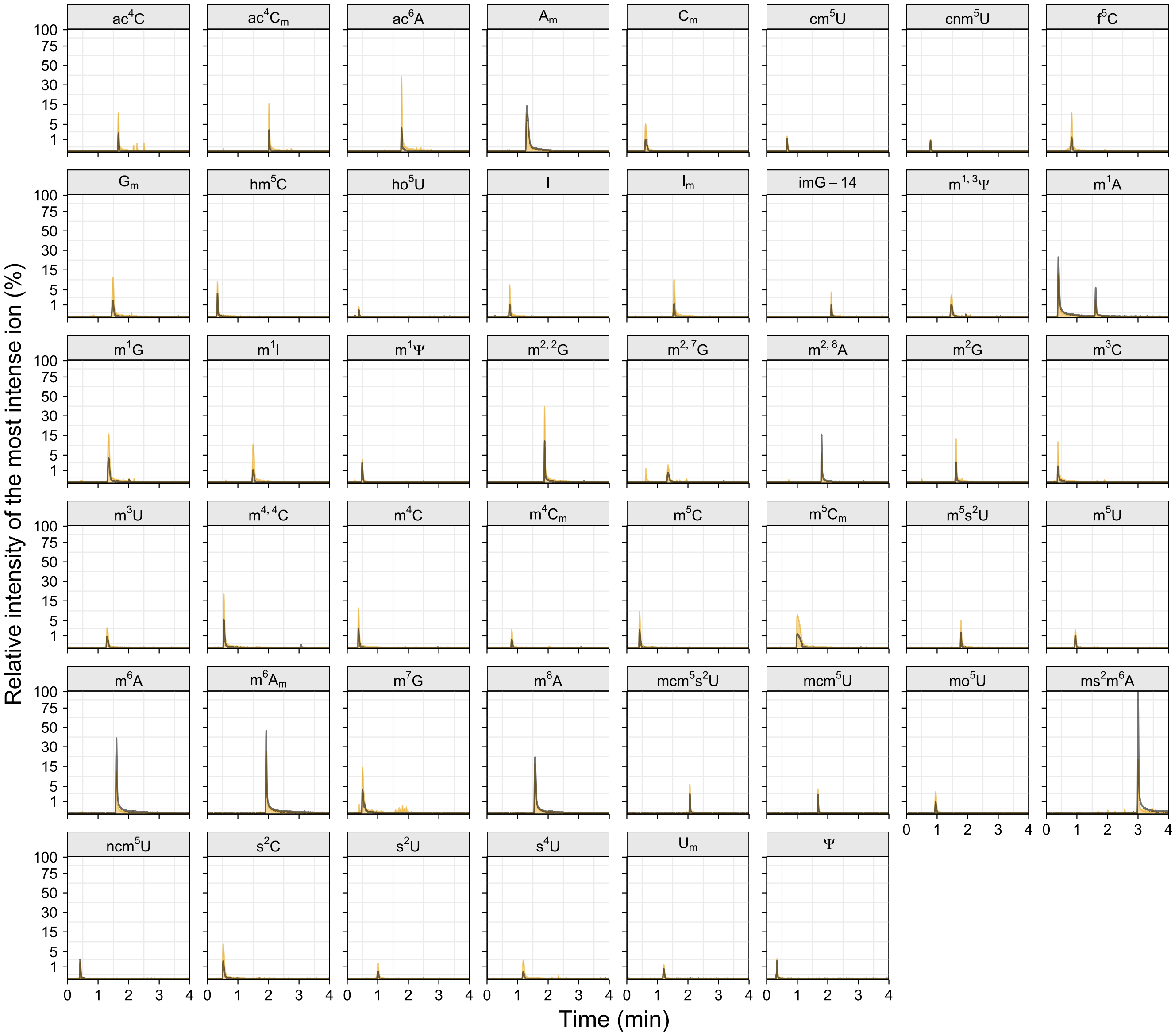


**Supplementary Figure S2.** Ionization propensity of the ribonucleoside standards on an ESI-QTOF instrument. Note the substantial difference between signal intensities of ribonucleosides presursor and product ions. Extracted ion chromatograms (XICs) of each modified ribonucleoside standard show that product ion (yellow) signals are, in some cases, more intense than precusor ions (black), as the sugar moiety (ribose) is easily fragmented from the mother ion (ribonucleoside) during ionization. To pinpoint the difference in signal intensities due to ionization efficiency, the relative intensity is calculated as proportion of the strongest detected signal, specifically $\mathbf{ms^2m^6A}$. The peak at ~1.5 min in $m^1A$ standard represents $m^6A$ formed by Dimroth rearrangement[1].

#### SUPPLEMENTARY FIGURE S3

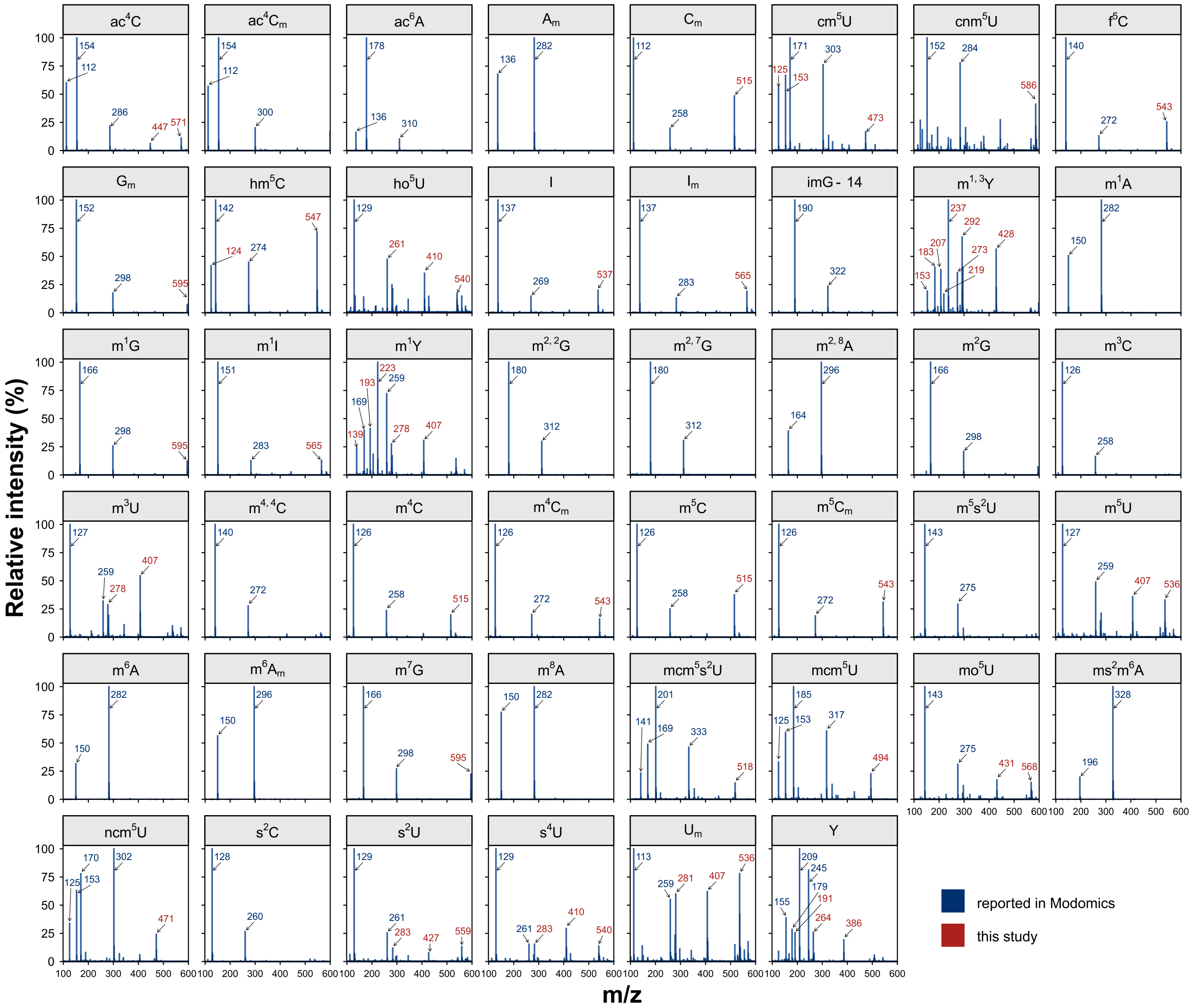


**Supplementary Figure S3.** Mass spectra of individual ribonucleoside standards detected by C18 UPLC-MS. The ribonucleoside indentity was confirmed by comparison or recorded spectra with previously reported ions (mass labelled in blue; reported in Modomics[2]). The additional recorded ions, specific to our system and ionization conditions are in red. Higher m/z ions represent adducts formed during ionization.

#### SUPPLEMENTARY FIGURE S4

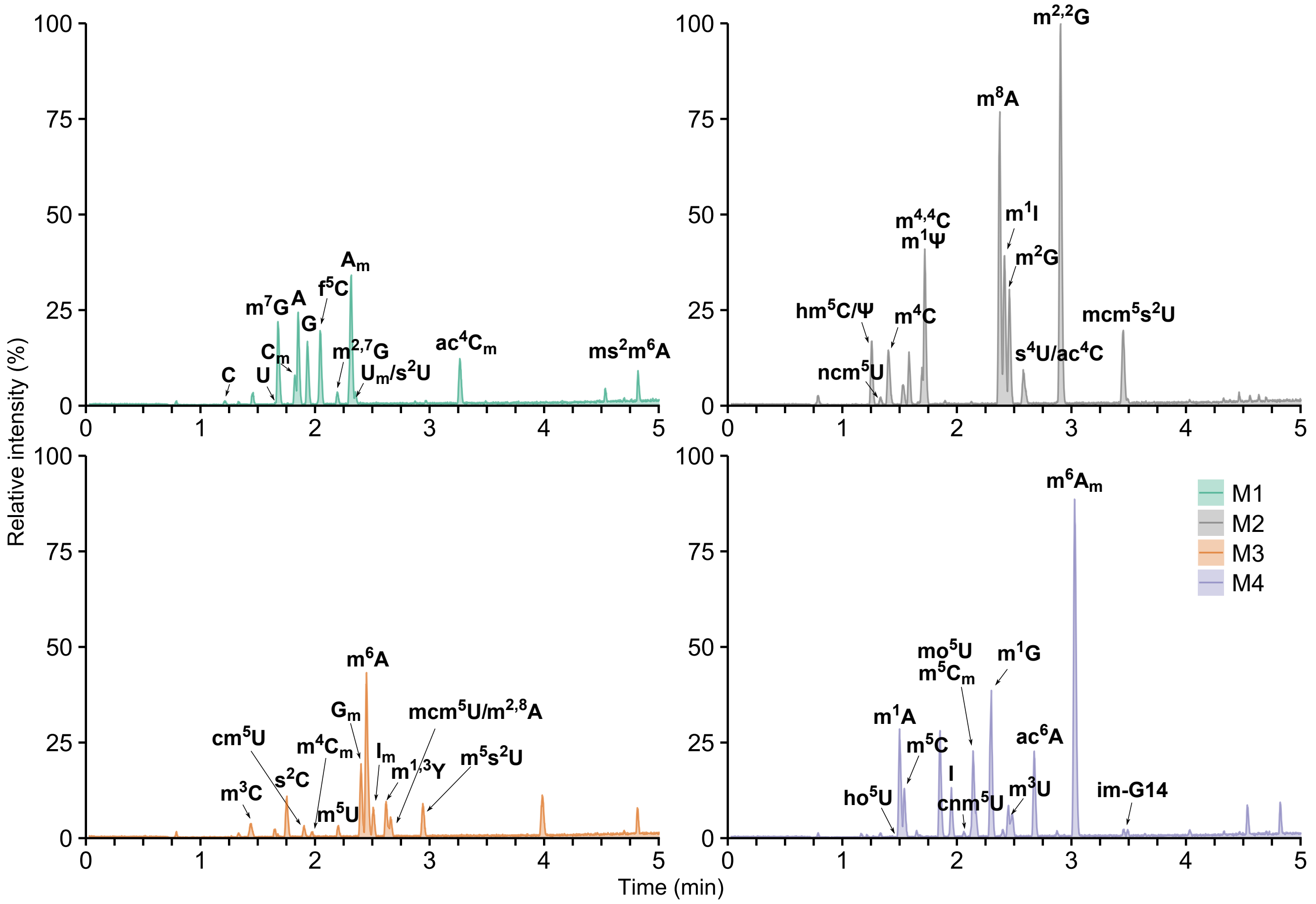


**Supplementary Figure S4.** Analysis of M1-M4 separation. Annotated TICs of four ribonucleoside standard mixes M1-M4.

#### SUPPLEMENTARY FIGURE S5

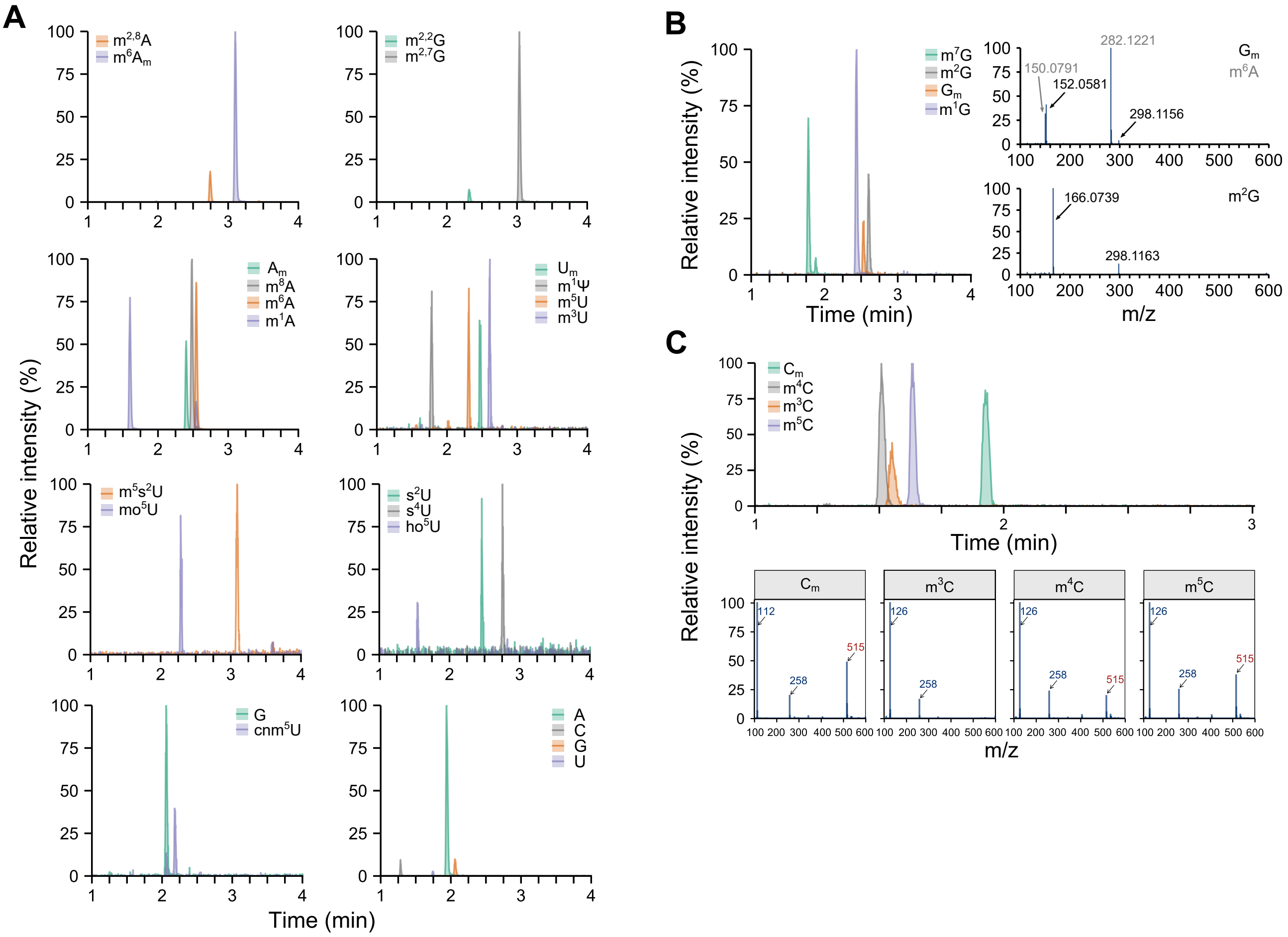


**Supplementary Figure S5.** Separation and identification of ribonucleoside isomers by C18 UPLC-MS. **(A)** Overlaid XICs showing the separation of isobaric ribonucleosides, positional isomers and canonical ribonucleosides. **(B)** XICs of selected positional isomers (left panel) and their respective MS spectra (right panel) showing the difference in ionization efficiencies. Neutral loss ion can be used as qualitative parameter to distinguish positional isomers. **(C)** Additional detected ions enhance the differentiation between positional isomers, which are difficult to separate such as methylated cytidines – $C_m$, $m^3C$, $m^4C$ and $m^5C$ (top panel). The dimer $[2M+H]^+$ ion 515 is not formed during ionization of $m^3C$ and thus may assist in discrimination between $m^3C$ and $m^4C$, which elute closely (bottom panel).

#### SUPPLEMENTARY FIGURE S6

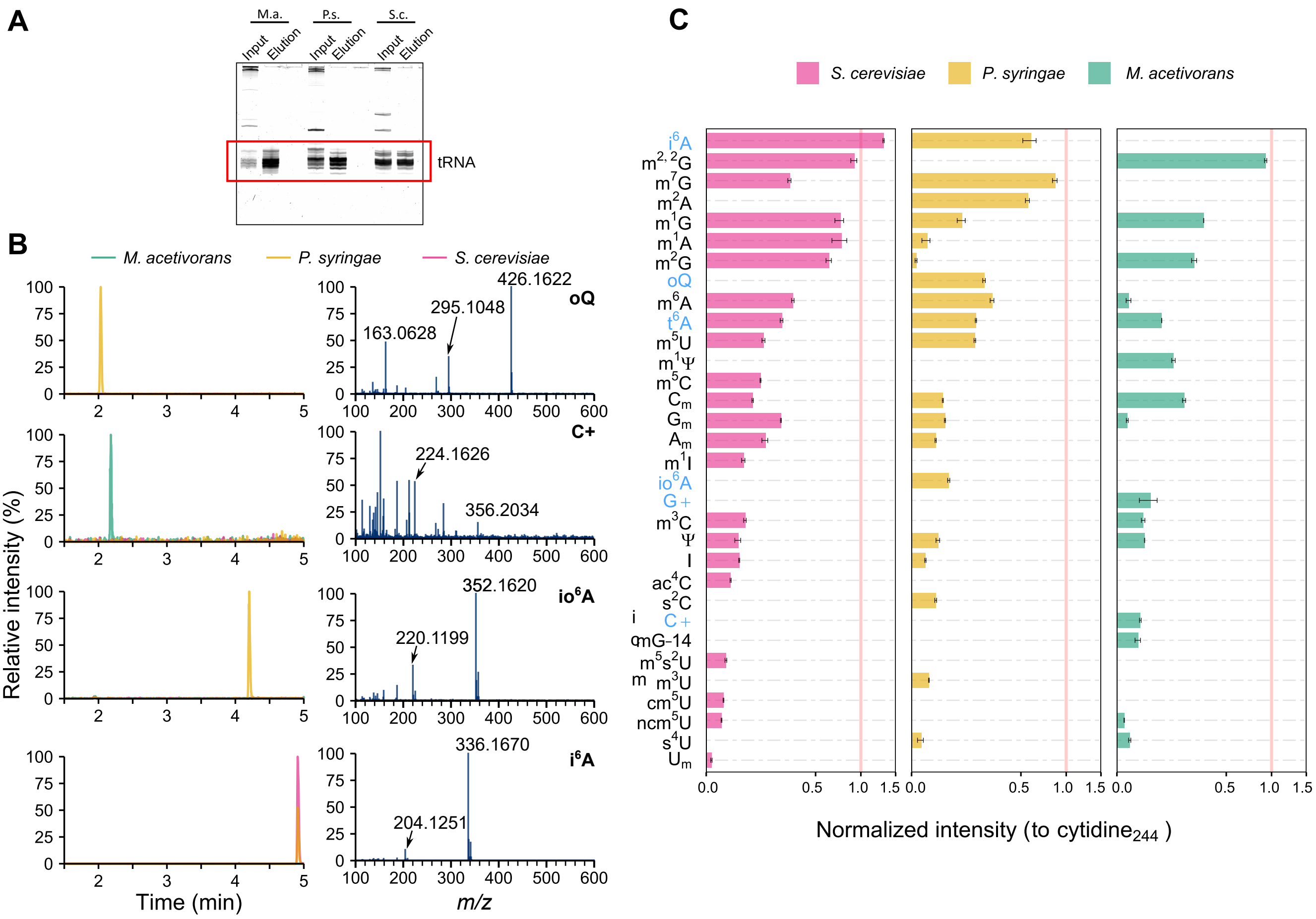


**Supplementary Figure S6.** Identification of unknown peaks present in biological samples and overview of identified modifications in studied microorganisms. **(A)** A representative denaturing polyacrylamide gel (10%) of total RNA (Input) and purified tRNA (Elution) isolated from all three microorganisms. **(B)** Overlaid XICs of all three organisms (left panel) and their respective mass spectra (right panel). Arrows point to neutral loss ions. **(C)** Identified ribonucleoside modifications in three representative organisms from each kingdom of life. Intensities are normalized to cytidine signal ($[M+H]^+$ ion 244). Modifications identified without standards are in blue. Error bars represent standard deviation (n = 3).

#### SUPPLEMENTARY FIGURE S7

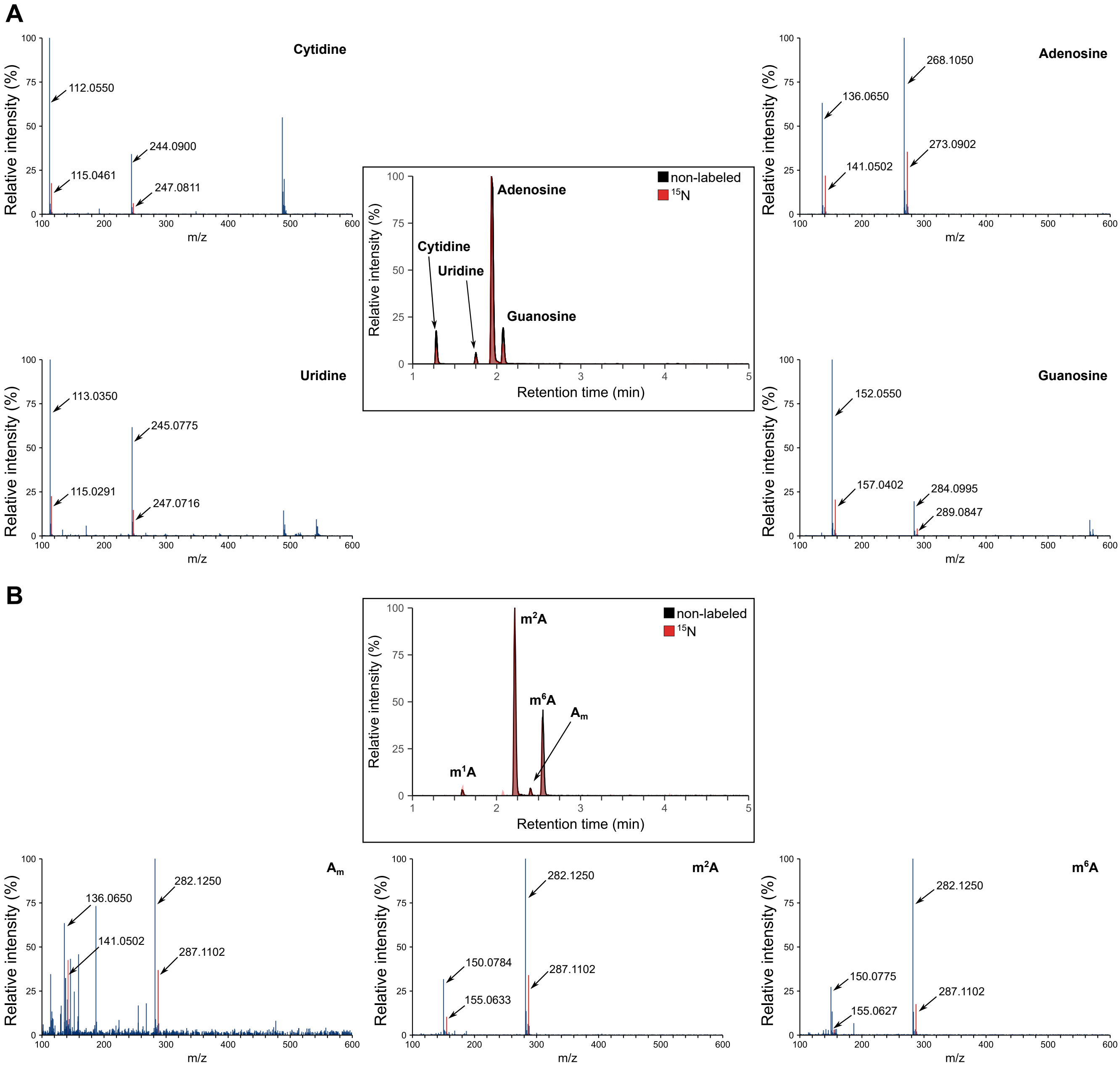


**Supplementary Figure S7.** Stable isotope ($^{15}$N) labeled ribonucleosides can be used as internal standard for ribonucleoside quantification. A mixture of non-labeled and $^{15}$N-labeled ribonucleosides, both originating from *Pseudomonas syringae*, was analyzed by C18 UPLC-MS. The overlays of XICs (central panel) show non-labeled **(A)** canonical ribonucleosides or **(B)** methylated adenosines and their $^{15}$N-labeled counterparts. The recorded spectra (side panels) show the mass difference between ions of non-labeled and $^{15}$N-labeled ribonucleosides.

### Supplementary Methods

#### Composition of ribonucleoside standard mixes

Each mix of standards consists of ribonucleosides mixed in equimolar amounts (100 nmol) and diluted to 10 µg/ml. Each mix (M1-M4) contains only one of the positional isomers to simplify their identification. The detailed composition of each mix is as follows:

**Supplementary Table S1: Detailed composition of Complete Mix (CM) and Limited Mix (LM; in grey)**

| Full name | Abbreviation | Monoisotopic mass | Amount in solution | | | | Purity* (%) |
|---|---|---|---|---|---|---|---|
| | | | nmol | µg | µg/ml (in LM) | µg/ml (in CM) | |
| adenosine | A | 267.0968 | 100 | 26.7 | 0.447 | 0.191 | 99 |
| uridine | U | 244.0695 | 100 | 24.4 | 0.409 | 0.175 | 100 |
| cytidine | C | 243.0855 | 100 | 24.3 | 0.407 | 0.174 | 99.9 |
| guanosine | G | 283.0917 | 100 | 28.3 | 0.474 | 0.202 | 98.7 |
| pseudouridine | Ψ | 244.0695 | 100 | 24.4 | 0.409 | 0.175 | 98.8 |
| 2'-O-methyladenosine | $A_m$ | 281.1124 | 100 | 28.1 | 0.471 | 0.201 | 99.9 |
| 2'-O-methyluridine | $U_m$ | 258.0852 | 100 | 25.8 | 0.432 | 0.185 | 99.8 |
| 2'-O-methylcytidine | $C_m$ | 257.1012 | 100 | 25.7 | 0.431 | 0.184 | 99.8 |
| 2'-O-methylguanosine | $G_m$ | 297.1073 | 100 | 28.7 | 0.481 | 0.205 | 99.9 |
| 2-thiouridine | $s^2U$ | 260.0467 | 100 | 26.0 | 0.436 | 0.186 | ≥98 |
| 4-thiouridine | $s^4U$ | 260.0467 | 100 | 26.0 | 0.436 | 0.186 | 98.9 |
| 5-methoxycarbonylmetyluridine | $mcm^5U$ | 316.0907 | 100 | 31.6 | 0.529 | 0.226 | 99.0 |
| *N4*-acetylcytidine | $ac^4C$ | 285.0961 | 100 | 28.5 | 0.478 | 0.204 | 98.9 |
| *N1*-methylguanosine | $m^1G$ | 297.1073 | 100 | 29.7 | 0.498 | 0.212 | 98.3 |
| *N2*-methylguanosine | $m^2G$ | 297.1073 | 100 | 29.7 | 0.498 | 0.212 | ≥95 |
| *N7*-methylguanosine | $m^7G$ | 297.1073 | 100 | 29.7 | 0.498 | 0.212 | ≥99.9 |
| *N1*-methyladenosine | $m^1A$ | 281.1124 | 100 | 28.1 | 0.471 | 0.201 | 99.8 |
| *N6*-methyladenosine | $m^6A$ | 281.1124 | 100 | 28.1 | 0.471 | 0.201 | 99.8 |
| *N3*-methylcytidine | $m^3C$ | 257.1012 | 100 | 25.7 | 0.431 | 0.184 | 97.2 |
| 5-methylcytidine | $m^5C$ | 257.1012 | 100 | 25.7 | 0.431 | 0.184 | 99.8 |
| *N3*-methyluridine | $m^3U$ | 258.0852 | 100 | 25.8 | 0.432 | 0.185 | ≥99.9 |
| 5-methyluridine | $m^5U$ | 258.0852 | 100 | 25.8 | 0.432 | 0.185 | 99.76 |
| *N6*-acetyladenosine | $ac^6A$ | 309.1073 | 100 | 30.9 | - | 0.221 | 98.0 |
| *N4*-acetyl-2'-O-methylcytidine | $ac^4C_m$ | 299.1117 | 100 | 29.9 | - | 0.214 | 99.5 |
| 5-carbamoylmethyluridine | $ncm^5U$ | 301.0910 | 100 | 30.1 | - | 0.215 | 98.1 |
| 5-(carboxymethyl)uridine | $cm^5U$ | 302.0750 | 100 | 30.2 | - | 0.216 | 96.2 |
| 5-(cyanomethyl)-uridine | $cnm^5U$ | 283.0804 | 100 | 28.3 | - | 0.202 | 90.0 |
| 4-demethylwyosine | imG-14 | 321.1073 | 100 | 32.1 | - | 0.230 | 96.4 |
| 2,8-dimethyladenosine | $m^{2,8}A$ | 295.1281 | 100 | 29.5 | - | 0.211 | 96.1 |
| *N4,N4*-dimethylcytidine | $m^{4,4}C$ | 271.1168 | 100 | 27.1 | - | 0.194 | 98.8 |
| *N2,N2*-dimethylguanosine | $m^{2,2}G$ | 311.1230 | 100 | 31.1 | - | 0.222 | 99.9 |
| *N2*,7-dimethylguanosine | $m^{2,7}G$ | 313.1386 | 100 | 31.2 | - | 0.223 | 95.1 |

*(Supplementary Table S1 continues on next page)*

*(Continued Supplementary Table S1)*

| | | | | | | | |
|---|---|---|---|---|---|---|---|
| 1,3-dimethylpseudouridine | $m^{1,3}\Psi$ | 272.1008 | 100 | 27.2 | - | 0.195 | 97.3 |
| 5-formylcytidine | $f^5C$ | 271.0804 | 100 | 27.1 | - | 0.194 | 99.3 |
| 5-hydroxymethylcytidine | $hm^5C$ | 273.0961 | 100 | 27.3 | - | 0.195 | 99.4 |
| 5-hydroxyuridine | $ho^5U$ | 260.0645 | 100 | 26.0 | - | 0.186 | 98.4 |
| inosine | I | 268.0808 | 100 | 26.8 | - | 0.192 | 99.6 |
| 5-methoxycarbonylmethyl-2-thiouridine | $mcm^5s^2U$ | 332.0678 | 100 | 33.2 | - | 0.238 | 98.8 |
| 5-methoxyuridine | $mo^5U$ | 274.0801 | 100 | 27.4 | - | 0.196 | 98.7 |
| 8-methyladenosine | $m^8A$ | 281.1124 | 100 | 28.1 | - | 0.201 | 99.9 |
| *N4*-methylcytidine | $m^4C$ | 257.1012 | 100 | 25.7 | - | 0.184 | >95 |
| *N1*-methylinosine | $m^1I$ | 282.0964 | 100 | 28.2 | - | 0.202 | 97.6 |
| 2’-O-methylinosine | $I_m$ | 282.0964 | 100 | 28.2 | - | 0.202 | 99 |
| *N6*-methyl-2’-O-methyladenosine | $m^6A_m$ | 295.1281 | 100 | 29.5 | - | 0.211 | 99.5 |
| *N4*-methyl-2’-O-methylcytidine | $m^4C_m$ | 271.1168 | 100 | 27.1 | - | 0.194 | 98.1 |
| 2’-O-methyl-5-methylcytidine | $m^5C_m$ | 271.1168 | 100 | 27.1 | - | 0.194 | 99.8 |
| 1-methylpseudouridine | $m^1\Psi$ | 258.0852 | 100 | 25.8 | - | 0.185 | 99.8 |
| 5-methyl-2-thiouridine | $m^5s^2U$ | 274.0623 | 100 | 27.4 | - | 0.196 | *n.a.* |
| 2-methylthio-*N6*-methyladenosine | $ms^2m^6A$ | 327.1001 | 100 | 32.7 | - | 0.234 | 98.4 |
| 2-thiocytidine | $s^2C$ | 259.0627 | 100 | 25.9 | - | 0.185 | 99.9 |
| | | **Final concentration of each mix**: | | | 10 | 10 | |

*as specified by manufacturer

**Supplementary Table S2: Composition of Ribonucleoside Mix 1 (M1)**

| Full name | Abbreviation | Monoisotopic mass | Amount in solution | | |
|---|---|---|---|---|---|
| | | | **nmol** | **µg** | **µg/ml** |
| cytidine | C | 243.0855 | 100 | 24.3 | 0.675 |
| uridine | U | 244.0695 | 100 | 24.4 | 0.678 |
| 2’-O-methylcytidine | $C_m$ | 257.1012 | 100 | 25.7 | 0.714 |
| 2’-O-methyluridine | $U_m$ | 258.0852 | 100 | 25.8 | 0.717 |
| 2-thiouridine | $s^2U$ | 260.0467 | 100 | 26.0 | 0.723 |
| adenosine | A | 267.0968 | 100 | 26.7 | 0.742 |
| 5-formylcytidine | $f^5C$ | 271.0804 | 100 | 27.1 | 0.753 |
| 2’-O-methyladenosine | $A_m$ | 281.1124 | 100 | 28.1 | 0.781 |
| guanosine | G | 283.0917 | 100 | 28.3 | 0.786 |
| *N7*-methylguanosine | $m^7G$ | 297.1073 | 100 | 29.7 | 0.825 |
| *N4*-acetyl-2’-O-methylcytidine | $ac^4C_m$ | 299.1117 | 100 | 29.9 | 0.831 |
| *N2,*7-dimethylguanosine | $m^{2,7}G$ | 313.1386 | 100 | 31.2 | 0.867 |
| 2-methylthio-*N6*-methyladenosine | $ms^2m^6A$ | 327.1001 | 100 | 32.7 | 0.909 |
| | | **Final concentration of each mix:** | | | 10 |

**Supplementary Table S3: Ribonucleoside mix 2 (M2)**

| Full name | Abbreviation | Monoisotopic mass | Amount in solution | | |
|---|---|---|---|---|---|
| | | | nmol | µg | µg/ml |
| pseudouridine | Ψ | 244.0695 | 100 | 27.3 | 0.748 |
| *N4*-methylcytidine | $m^4C$ | 257.1012 | 100 | 24.4 | 0.668 |
| 1-methylpseudouridine | $m^1\Psi$ | 258.0852 | 100 | 25.7 | 0.704 |
| 4-thiouridine | $s^4U$ | 260.0467 | 100 | 30.1 | 0.824 |
| *N4,N4*-dimethylcytidine | $m^{4,4}C$ | 271.1168 | 100 | 25.8 | 0.706 |
| 5-hydroxymethylcytidine | $hm^5C$ | 273.0961 | 100 | 27.1 | 0.742 |
| 8-methyladenosine | $m^8A$ | 281.1124 | 100 | 26.0 | 0.712 |
| *N1*-methylinosine | $m^1I$ | 282.0964 | 100 | 28.2 | 0.772 |
| *N4*-acetylcytidine | $ac^4C$ | 285.0961 | 100 | 28.1 | 0.769 |
| *N2*-methylguanosine | $m^2G$ | 297.1073 | 100 | 29.7 | 0.813 |
| 5-carbamoylmethyluridine | $ncm^5U$ | 301.0910 | 100 | 28.5 | 0.780 |
| *N2,N2*-dimethylguanosine | $m^{2,2}G$ | 311.1230 | 100 | 31.1 | 0.852 |
| 5-methoxycarbonylmethyl-2-thiouridine | $mcm^5s^2U$ | 332.0678 | 100 | 33.2 | 0.909 |
| | | **Final concentration of each mix:** | | | 10 |

**Supplementary Table S4: Ribonucleoside mix 3 (M3)**

| Full name | Abbreviation | Monoisotopic mass | Amount in solution | | |
|---|---|---|---|---|---|
| | | | nmol | µg | µg/ml |
| *N3*-methylcytidine | $m^3C$ | 257.1012 | 100 | 25.7 | 0.383 |
| 5-methyluridine | $m^5U$ | 258.0852 | 100 | 25.8 | 0.385 |
| 2-thiocytidine | $s^2C$ | 259.0627 | 100 | 25.9 | 0.386 |
| *N4*-methyl-2'-O-methylcytidine | $m^4C_m$ | 271.1168 | 100 | 27.1 | 0.404 |
| 1,3-dimethylpseudouridine | $m^{1,3}\Psi$ | 272.1008 | 100 | 27.2 | 0.405 |
| 5-methyl-2-thiouridine | $m^5s^2U$ | 274.0623 | 100 | 27.4 | 0.409 |
| *N6*-methyladenosine | $m^6A$ | 281.1124 | 100 | 28.1 | 0.419 |
| 2'-O-methylinosine | $I_m$ | 282.0964 | 100 | 28.2 | 0.420 |
| 2,8-dimethyladenosine | $m^{2,8}A$ | 295.1281 | 100 | 29.5 | 0.440 |
| 2'-O-methylguanosine | $G_m$ | 297.1073 | 100 | 28.7 | 0.428 |
| 5-(carboxymethyl)uridine | $cm^5U$ | 302.0750 | 100 | 30.2 | 0.450 |
| 5-methylcarbonylmetyluridine | $mcm^5U$ | 316.0907 | 100 | 31.6 | 0.471 |
| | | **Final concentration of each mix:** | | | 10 |

**Supplementary Table S5: Ribonucleoside mix 4 (M4)**

| Full name | Abbreviation | Monoisotopic mass | Amount in solution | | |
|---|---|---|---|---|---|
| | | | nmol | µg | µg/ml |
| 5-methylcytidine | $m^5C$ | 257.1012 | 100 | 25.7 | 0.762 |
| *N3*-methyluridine | $m^3U$ | 258.0852 | 100 | 25.8 | 0.765 |
| 5-hydroxyuridine | $ho^5U$ | 260.0645 | 100 | 26.0 | 0.770 |
| inosine | I | 268.0808 | 100 | 26.8 | 0.794 |
| 2'-O-methyl-5-methylcytidine | $m^5C_m$ | 271.1168 | 100 | 27.1 | 0.803 |
| 5-methoxyuridine | $mo^5U$ | 274.0801 | 100 | 27.4 | 0.812 |
| *N1*-methyladenosine | $m^1A$ | 281.1124 | 100 | 28.1 | 0.833 |
| 5-(cyanomethyl)-uridine | $cnm^5U$ | 283.0804 | 100 | 28.3 | 0.839 |
| *N6*-methyl-2'-O-methyladenosine | $m^6A_m$ | 295.1281 | 100 | 29.5 | 0.874 |
| *N1*-methylguanosine | $m^1G$ | 297.1073 | 100 | 29.7 | 0.880 |
| *N6*-acetyladenosine | $ac^6A$ | 309.1073 | 100 | 30.9 | 0.916 |
| 4-demethylwyosine | imG-14 | 321.1073 | 100 | 32.1 | 0.951 |
| | | **Final concentration of each mix:** | | | 10 |

**Cultivation medium for *Methanosarcina acetivorans***

The high-salt broth medium used for cultivation of *M. acetivorans* consists of the following components: 400 mM NaCl, 45 mM $NaHCO_3$, 13 mM KCl, 0.00005% (w/v) resazurine, trace element solution (8 mM nitrilotriacetic acid, 12 mM $MgSO_4$, 3 mM $MnSO_4$, 17 mM NaCl, 0.36 mM $FeSO_4$, 0.64 mM $CoSO_4$, 0.68 mM $CaCl_2$, 0.63 mM $ZnSO_4$, 40 µM $CuSO_4$, 42 µM $KAl(SO_4)_2$, 0.16 mM $H_3BO_3$, 41 µM $Na_2MoO_4$, 0.13 mM $NiCl_2$, 1 µM $Na_2SeO_3$, 1.2 µM $Na_2WO_4$), vitamin solution (2 mg $L^{-1}$ biotin, 2 mg/l folic acid, 10 mg/l pyridoxine-HCl, 5 mg/l thiamine-HCl, 5 mg/l riboflavin, 5 mg/l nicotinic acid, 5 mg/l D-Ca-pantothenate, 0.1 mg/l vitamin $B_{12}$, 5 mg/l p-aminobenzoic acid, 5 mg/l lipoic acid, 54 mM $MgCl_2$, 2 mM $CaCl_2$).

**UPLC method development for ribonucleoside separation**

We explored the suitability of three different stationary phases for baseline separation of modified ribonucleosides. To this end, we tested the following columns – Acquity UPLC® BEH HILIC (1.7 µm, 2.1 × 100 mm, Waters, Ireland), Acquity UPLC® BEH Amide (1.7 µm, 2.1 × 100 mm, Waters, Ireland), Acquity UPLC® BEH C18 (1.7 µm, 2.1 × 50 mm and 2.1 × 150 mm, Waters, Ireland) – with varying chromatographic parameters (temperature, flow rate, gradient, solvent pairs, and solvent pH) and ionization modes (ESI+/-) as outlined below:

I) Acquity UPLC® BEH Amide column with solvents, **A**: 10 mM ammonium bicarbonate, pH 9.0; **B**: ACN (ESI-)

II) Acquity UPLC® BEH Amide column with solvents, **A**: 10 mM ammonium bicarbonate, pH 9.0; **B**: ACN (ESI+)

III) Acquity UPLC® BEH Amide column with solvents, **A**: 0.1% formic acid in $H_2O$ **B**: 0.1% formic acid in ACN (ESI+)

IV) Acquity UPLC® BEH HILIC column with solvents, **A**: 0.1% formic acid in $H_2O$ **B**: 0.1% formic acid in ACN (ESI+)

V) Acquity UPLC® BEH C18 columns (both 50 and 150 mm) with solvents, **A**: 0.1% formic acid in MQ $H_2O$ **B**: 0.1% formic acid in acetonitrile (ESI+)

All UPLC-MS analyses were executed with Waters Acquity UPLC® system (Waters, Milford MA, USA) attached to Waters Synapt G2 HDMS mass spectrometer (Waters, Milford MA, USA) via an ESI ion source. Samples were analyzed in negative (I) and/or positive (II-V) sensitivity ion mode. The mass/charge (*m/z*) range was set from 100 to 600. The separation of modified ribonucleosides was tested using the abovementioned (I-V) combinations of solvents and columns.

Briefly, the separation on Aquity UPLC® BEH Amide column and Acquity UPLC® BEH HILIC column (both 1.7 µm, 2.1 × 100 mm, Waters, Ireland) was performed at +35°C with solvents as noted above. A linear gradient started at 5% of A, was held for 1 min, then increased to 35% in 5 min, to 65% in 6 min, switched back to 5% and finally left to stabilize for 2 min with a total run time of 8 min. The flow rate of the mobile phase was 0.4 ml/min, sample loading 50 ng and tray temperature was set to +12 °C. Capillary voltage was set to at 1.5 kV, sampling cone 40 and extraction cone 4.0, the source temperature 120 °C and desolvation temperature 400 °C, cone and desolvation gas flow rate was set to 20 l/h and 800 l/h, respectively.

The detailed conditions of ribonucleosides separation on Acquity UPLC® BEH C18 columns are described in the Experimental section.

#### Calculation of chromatographic parameters

Chromatographic parameters (Supplementary Table S6) were calculated as following:

Tailing factor $TF = (f + b)/2f$, and

Peak asymmetry $A_s = b/f$,

where *f* and *b* is front and back peak width at 5% for TF and 10% for $A_s$ of the maximum intensity, respectively.

Theoretical number of plates $N = 5.54 * (t_R/W_{0.5})^2$,

where $t_R$ is retention time at peak apex and $W_{0.5}$ is peak Full Width at Half Maximum (FWHM). FWHM was calculated in R with IDPmisc package (version 1.1.20).

**Supplementary Table S6: Chromatographic properties of 50 mm Acquity UPLC® BEH C18 column for the separation of modified ribonucleosides.**

| Ribonucleoside Abbreviation | Full Width at Half Maximum (FWHM) ($\times 10^{-2}$ min) | Tailing Factor (TF) | Peak Asymmetry (As) | Theoretical Number of Plates (N) |
|---|---|---|---|---|
| Ψ | 1.45 | 0.88 | 1.14 | 3058 |
| $A_m$ | 5.59 | 1.91 | 2.53 | 3026 |
| $U_m$ | 2.90 | 1.30 | 1.57 | 9670 |
| $C_m$ | 4.14 | 2.45 | 3.78 | 1200 |
| $G_m$ | 5.06 | 1.17 | 1.25 | 4775 |
| $s^2U$ | 3.11 | 1.25 | 1.39 | 5843 |
| $s^4U$ | 2.90 | 1.32 | 1.61 | 9329 |
| $mcm^5U$ | 2.07 | 1.31 | 1.57 | 36359 |
| $ac^4C$ | 1.86 | 1.28 | 1.37 | 44327 |
| $m^1G$ | 4.14 | 1.32 | 1.60 | 5854 |
| $m^2G$ | 3.40 | 1.02 | 1.50 | 12537 |
| $m^7G$ | 3.31 | 2.87 | 3.83 | 1275 |
| $m^1A$ | 4.23 | 1.50 | 3.29 | 478 |
| $m^6A$ | 1.45 | 1.26 | 1.31 | 67909 |
| $m^3C$ | 2.58 | 2.16 | 2.69 | 1196 |
| $m^5C$ | 2.48 | 2.29 | 3.00 | 1537 |
| $m^3U$ | 4.65 | 1.54 | 1.83 | 4322 |
| $m^5U$ | 2.48 | 1.32 | 1.60 | 8081 |
| $ac^6A$ | 2.78 | 1.39 | 1.42 | 22698 |
| $ac^4C_m$ | 1.66 | 1.36 | 1.43 | 82694 |
| $ncm^5U$ | 2.07 | 1.64 | 1.86 | 2234 |
| $cm^5U$ | 2.28 | 1.56 | 1.88 | 4696 |
| $cnm^5U$ | 2.07 | 1.35 | 1.56 | 7987 |
| imG-14 | 0.62 | 1.70 | 2.49 | 643740 |
| $m^{2,8}A$ | 2.99 | 1.57 | 2.00 | 19984 |
| $m^{4,4}C$ | 1.24 | 1.84 | 2.30 | 10475 |
| $m^{2,2}G$ | 1.86 | 1.32 | 1.58 | 56818 |
| $m^{2,7}G$ | 4.55 | 1.75 | 2.46 | 4811 |
| $m^{1,3}\Psi$ | 5.27 | 1.41 | 1.93 | 4297 |
| $f^5C$ | 2.48 | 1.46 | 1.55 | 6119 |
| $hm^5C$ | 1.86 | 1.58 | 2.20 | 1738 |
| $ho^5U$ | 2.37 | 1.37 | 1.48 | 1428 |
| I | 3.61 | 1.26 | 1.29 | 2330 |
| $mcm^5s^2U$ | 1.86 | 1.29 | 1.43 | 67757 |
| $mo^5U$ | 2.69 | 1.15 | 1.27 | 6969 |
| $m^8A$ | 2.91 | 1.31 | 1.39 | 16277 |
| $m^4C$ | 1.04 | 2.45 | 3.89 | 6898 |
| $m^1I$ | 3.52 | 1.11 | 1.13 | 10152 |
| $I_m$ | 2.69 | 1.03 | 0.92 | 18385 |
| $m^6A_m$ | 1.86 | 1.36 | 1.27 | 59472 |
| $m^4C_m$ | 3.11 | 1.36 | 1.70 | 3780 |

*(Supplementary Table S6 continues on next page)*

(Continued Supplementary Table S6)

| | | | | |
|---|---|---|---|---|
| $m^5C_m$ | 7.89 | 2.74 | 4.57 | 902 |
| $m^1\Psi$ | 1.86 | 1.79 | 1.86 | 3834 |
| $m^5s^2U$ | 1.86 | 1.19 | 1.16 | 50757 |
| $ms^2m^6A$ | 1.66 | 1.35 | 1.54 | 182645 |
| $s^2C$ | 2.90 | 2.03 | 2.93 | 1778 |

**Multiple Reaction Monitoring (MRM) using QQQ detector**

For high-sensitivity and targeted ribonucleoside detection, we coupled Acquity UPLC® BEH C18 column (Ø 1.7 µm, 2.1 × 150 mm, Waters, Ireland) to ExionLC UPLC with ABSciex QTRAP-6500+ detector and analyzed ribonucleoside standards by multiple reaction monitoring (MRM) using ESI+ mode. The transitions and other parameters for tested ribonucleoside standards the following used in MRM method are noted in Supplementary Table S7.

**Supplementary Table S7: Parameters used for multiple-reaction monitoring (MRM)**

| Ribonucleoside Abbreviation | Precursor m/z | Product m/z | Declustering Potential (DP, volts) | Entrance potential (EP, volts) | Collision energy (CE, volts) | Collision Cell Exit Potential (CXP, volts) | Retention time (min) |
|---|---|---|---|---|---|---|---|
| C | 244.090 | 112.050 | 5 | 10.0 | 45.0 | 12.0 | 1.25 |
| U | 245.080 | 113.040 | 5 | 10.0 | 17.0 | 12.0 | 1.71 |
| Ψ | 245.080 | 155.060 | 5 | 10.0 | 15.0 | 14.0 | 1.25 |
| $m^4C$ | 258.100 | 126.070 | 5 | 10.0 | 21.0 | 14.0 | 1.45 |
| $m^3C$ | 258.100 | 126.050 | 5 | 10.0 | 35.0 | 14.0 | 2.00 |
| $m^5C$ | 258.100 | 126.060 | 5 | 10.0 | 17.0 | 14.0 | 1.57 |
| $C_m$ | 258.110 | 112.050 | 5 | 10.0 | 25.0 | 14.0 | 1.63 |
| $U_m$ | 259.090 | 113.040 | 5 | 10.0 | 17.0 | 14.0 | 2.40 |
| $m^5U$ | 259.090 | 127.050 | 5 | 10.0 | 19.0 | 14.0 | 2.30 |
| $m^3U$ | 259.090 | 127.050 | 5 | 10.0 | 17.0 | 16.0 | 2.57 |
| $m^1\Psi$ | 259.100 | 169.060 | 5 | 10.0 | 17.0 | 10.0 | 1.76 |
| $s^2C$ | 260.070 | 128.050 | 5 | 10.0 | 23.0 | 14.0 | 1.87 |
| $s^2U$ | 261.050 | 129.030 | 5 | 10.0 | 15.0 | 6.0 | 2.40 |
| $s^4U$ | 261.050 | 129.010 | 5 | 10.0 | 17.0 | 14.0 | 2.67 |
| $ho^5U$ | 261.070 | 129.010 | 5 | 10.0 | 25.0 | 18.0 | 1.50 |
| A | 268.100 | 136.060 | 5 | 10.0 | 23.0 | 12.0 | 2.00 |
| I | 269.090 | 137.050 | 5 | 10.0 | 21.0 | 14.0 | 2.04 |
| $f^5C$ | 272.090 | 140.050 | 5 | 10.0 | 17.0 | 14.0 | 2.15 |
| $m^4C_m$ | 272.120 | 126.050 | 5 | 10.0 | 17.0 | 20.0 | 2.10 |
| $m^5C_m$ | 272.120 | 126.070 | 5 | 10.0 | 21.0 | 12.0 | 2.26 |
| $m^{4,4}C$ | 272.130 | 140.080 | 5 | 10.0 | 17.0 | 4.0 | 1.80 |
| $m^{1,3}\Psi$ | 273.100 | 153.050 | 5 | 10.0 | 25.0 | 10.0 | 2.50 |
| $hm^5C$ | 274.100 | 142.060 | 5 | 10.0 | 27.0 | 6.0 | 1.25 |

(Supplementary Table S7 continues on next page)

*(Continued Supplementary Table S7)*

| | | | | | | | |
|---|---|---|---|---|---|---|---|
| $m^5s^2U$ | 275.070 | 143.050 | 5 | 10.0 | 15.0 | 28.0 | 3.05 |
| $mo^5U$ | 275.090 | 143.050 | 5 | 10.0 | 17.0 | 18.0 | 2.25 |
| $A_m$ | 282.120 | 136.060 | 5 | 10.0 | 21.0 | 14.0 | 2.40 |
| $m^8A$ | 282.120 | 150.080 | 5 | 10.0 | 31.0 | 18.0 | 2.48 |
| $m^6A$ | 282.120 | 150.050 | 5 | 10.0 | 25.0 | 18.0 | 2.55 |
| $m^1A$ | 282.120 | 150.080 | 5 | 10.0 | 29.0 | 20.0 | 1.58 |
| $m^1I$ | 283.100 | 151.060 | 5 | 10.0 | 21.0 | 14.0 | 2.50 |
| $I_m$ | 283.100 | 137.050 | 5 | 10.0 | 29.0 | 14.0 | 2.55 |
| $cnm^5U$ | 284.090 | 152.050 | 5 | 10.0 | 27.0 | 20.0 | 2.15 |
| G | 284.100 | 152.050 | 5 | 10.0 | 19.0 | 14.0 | 2.00 |
| $ac^4C$ | 286.100 | 154.060 | 5 | 10.0 | 17.0 | 10.0 | 2.69 |
| $m^{2,8}A$ | 296.140 | 164.050 | 5 | 10.0 | 29.0 | 14.0 | 2.75 |
| $m^6A_m$ | 296.140 | 150.080 | 5 | 10.0 | 21.0 | 18.0 | 3.13 |
| $G_m$ | 298.100 | 152.050 | 5 | 10.0 | 21.0 | 8.0 | 2.51 |
| $m^7G$ | 298.120 | 166.070 | 5 | 10.0 | 17.0 | 10.0 | 1.80 |
| $m^2G$ | 298.120 | 166.070 | 5 | 10.0 | 21.0 | 18.0 | 2.56 |
| $m^1G$ | 298.120 | 166.070 | 5 | 10.0 | 21.0 | 20.0 | 2.40 |
| $ac^4C_m$ | 300.120 | 154.060 | 5 | 10.0 | 13.0 | 8.0 | 3.40 |
| $ncm^5U$ | 302.100 | 170.050 | 5 | 10.0 | 15.0 | 16.0 | 1.60 |
| $cm^5U$ | 303.080 | 171.050 | 5 | 10.0 | 15.0 | 20.0 | 2.00 |
| $ac^6A$ | 310.120 | 178.070 | 5 | 10.0 | 17.0 | 14.0 | 2.77 |
| $m^{2,7}G$ | 312.130 | 180.060 | 5 | 10.0 | 25.0 | 26.0 | 2.30 |
| $m^{2,2}G$ | 312.130 | 180.080 | 5 | 10.0 | 19.0 | 12.0 | 3.00 |
| $mcm^5U$ | 317.100 | 153.050 | 5 | 10.0 | 29.0 | 16.0 | 2.75 |
| imG-14 | 322.120 | 190.070 | 5 | 10.0 | 25.0 | 16.0 | 3.56 |
| $ms^2m^6A$ | 328.110 | 196.070 | 5 | 10.0 | 27.0 | 10.0 | 4.90 |
| $mcm^5s^2U$ | 333.070 | 201.030 | 5 | 10.0 | 17.0 | 14.0 | 3.54 |

#### References


1. Macon JB, Wolfenden R. 1-Methyladenosine. Dimroth rearrangement and reversible reduction. Biochemistry 1968; 7:3453–8. https://doi.org/10.1021/bi00850a021

2. Boccaletto P, Machnicka MA, Purta E, Piatkowski P, Baginski B, Wirecki TK, de Crécy-Lagard V, Ross R, Limbach PA, Kotter A, et al. MODOMICS: a database of RNA modification pathways. 2017 update. Nucleic Acids Res 2018; 46:D303–7. PMID:29106616; https://doi.org/10.1093/nar/gkx1030
